# Durotaxis bridges phase transition as a function of tissue stiffness *in vivo*

**DOI:** 10.1101/2023.01.05.522913

**Authors:** Min Zhu, Bin Gu, Evan Thomas, Hirotaka Tao, Theodora M. Yung, Kaiwen Zhang, Janet Rossant, Yu Sun, Sevan Hopyan

**Author notes:** Authors for correspondence: Sevan Hopyan, 686 Bay Street, 16-9713, Toronto, Ontario M5G 0A4, Canada, or Yu Sun, 5 King’s College Road, MC419, Toronto, Ontario M5S 3G8.

## Abstract

Physical processes ultimately drive morphogenetic cell movements. Two proposals are that 1) cells migrate toward stiffer tissue (durotaxis) and that 2) the extent of cell rearrangements reflects liquid-solid tissue phase. It is unclear whether and how these concepts are related. Here, we identify fibronectin-dependent tissue stiffness as a control variable that underlies and unifies these phenomena *in vivo*. In murine limb bud mesoderm, cells are either caged, move directionally by durotaxis or intercalate as a function of their location along a stiffness gradient. A unifying stiffness-phase transition model that is based on a Landau equation accurately predicts cell diffusivity upon loss or gain of fibronectin. Fibronectin is regulated by a WNT5A-YAP positive feedback pathway that controls cell movements, tissue shape and skeletal pattern. The results identify a key determinant of phase transition and show how durotaxis emerges in a mixed phase environment *in vivo*.

## INTRODUCTION

Although cell division underlies growth, morphogenesis involves fundamentally physical processes that form the embryo. Cell movements, rather than the spatial distributions of cell divisions, underlie the shapes of various embryonic structures^1,2^. Long-range coordination of morphogenetic cell movements shapes embryonic structures such as the main body axis^3^, the face^2^, limbs^4^, skin^5^, and other organ primordia^6^. Biochemical cues such as morphogens, chemoattractants and cell polarity pathways^6,7^ do not fully explain long-range cell coordination *in vivo*, in part because of the short range^8^ and noisy nature^9^ of diffusible ligand gradients. Physical cues such as tissue stiffness represent strong candidate mechanisms because they can be rapidly communicated among hundreds or thousands of cells^4,10^. Likely a combination of mechanisms fully explains morphogenesis^11^.

Liquid-solid phase transition is a relatively recent and potentially useful physical framework to describe cell and tissue movements^12,13^. The concept is based, in part, on the physics of foams to explain the frequency of cell rearrangements and has been assessed as a function of cellular parameters^3,12,14,15^. This paradigm has been applied to describe elongation of the zebrafish tail bud^3^, vertebrate digit primordia^16^, and the mandibular arch of the mouse embryo^4^. In the theoretical case of confluent cells, tissue phase can be determined by cell geometries^14^ although most tissues are not confluent. Jamming is another type of phase transition in which cell packing correlates with structural tissue integrity and unjammed cells rearrange in a manner that morphs tissue. However, mechanistic explanations for this phenomenon are incomplete. Jamming correlates with the degree of cell-cell proximity (connectivity)^12^ or adhesion^3^ in zebrafish. In contrast to particulate (multicellular)^12^ or continuum (bulk tissue)^3^ models that have been used to describe phase transition, tissues typically contain both cells and substrates that profoundly impact material properties. The contribution of tissue substrate to tissue phase has not been studied. Moreover, although tissue stiffness corresponds to cell jamming^3^, the relationship between material tissue properties and tissue phase remains unclear.

Durotaxis is another, distinct, hypothesis that has been used to explain the directional movement of cells. As has been shown primarily *in vitro*^17^, cells preferentially move toward stiffer substrate. Evidence for durotaxis *in vivo* was shown during neural crest cell migration in *Xenopus*^11,18^ and in correlative findings in our previous work on the mouse limb bud^4^. However, the relationship between material tissue properties and the orientation of cell movements remains unclear. Experiments that supported durotaxis in *Xenopus* involved epithelial dissection and exposed-surface indentation methods to measure subsurface stiffness gradients, and manipulation of stiffness by mechanical means *in vivo* and perturbation *in vitro*, thereby introducing confounding variables such as altered tissue integrity and uncertain cellular resolution *in vivo*^11,18^. Nonetheless, durotaxis is an important consideration, especially in a mixed phase environment in which a stiffness gradient would be expected to interpolate between liquid and solid states, a possibility that has not been explored.

Here, we manipulated tissue stiffness *in vivo* by performing conditional loss and gain of function experiments using a floxed fibronectin (*Fn*) and a knock-in Fn overexpression strain that we generated. Using live light sheet time-lapse imaging of embryos harbouring a transgenic far-red nuclear reporter (H2B-miRFP703^19^), we identified a range of cell movement types that vary as a function of FN-dependent stiffness. Cells in the stiffest and softest regions of limb bud mesoderm exhibited caged or rearranging behaviours, respectively, with low directional persistence (a measure of directional constancy). In the intervening region where a stiffness gradient dominates, cells moved in a coordinated, durotactic manner with high persistence. Starting with a Landau theory for second order phase transitions, we constructed a model with a term describing the coupling of cellular diffusion to a stiffness gradient. This stiffness-phase transition model accurately predicted cell diffusivity as a function of stiffness upon loss and gain of fibronectin *in vivo*. Using additional mutants, we showed that FN is downstream of a WNT5A-YAP pathway and feeds-back to promote YAP nuclear translocation and *Wnt5a* expression. The results identify extracellular matrix abundance as a determinant of phase transition and show how durotaxis emerges in a mixed phase environment *in vivo*. They demonstrate how stiffness affects tissue shape and subsequent skeletal pattern.

## RESULTS

### Modes of mesodermal cell movements vary as a function of tissue stiffness

To obtain a 3D tissue stiffness map, we utilised a 3D magnetic tweezer system that we developed previously^4^. By generating a uniform magnetic field gradient within the work-space, magnetic beads injected in the E9.25 mouse limb bud were actuated with identical forces (Figure 1A). Local tissue stiffness was quantified by fitting the bead displacement-time curve using a Zener model with a serial dashpot. To merge tissue stiffness maps obtained from different embryos while avoiding data skewing by sub-somite staging differences in limb bud morphologies, we performed 3D limb bud shape registration using a customised program^20^ (Methods, Figures S1A–S1C and Movie S1). As before^4^, we confirmed that the 21 somite (som., ∼E9.25) mouse limb bud exhibited an anteroproximally biased mesodermal stiffness gradient (Figures 1B and S1D).

**Figure 1.**
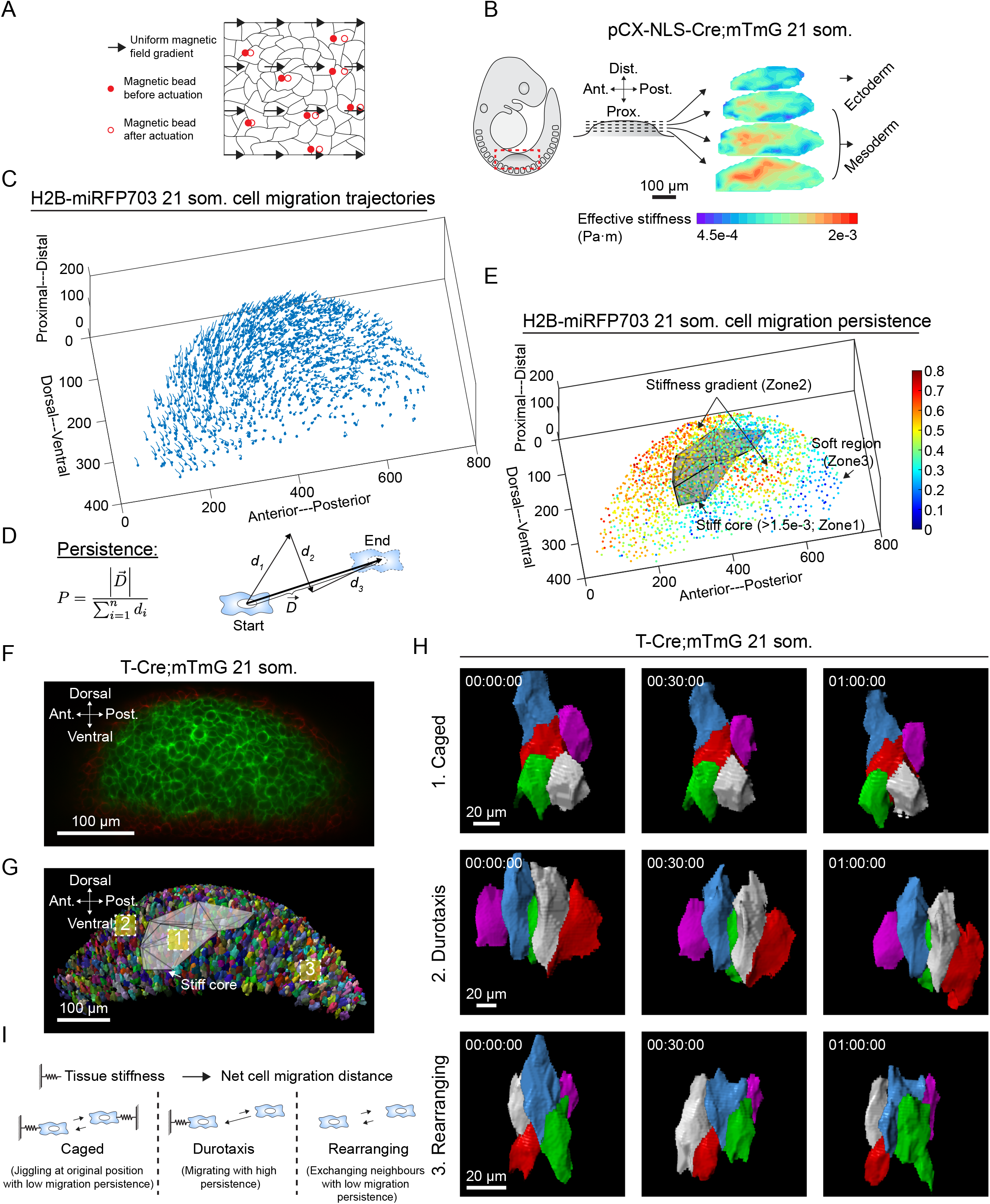
Zonal cell movements correspond to three-dimensional tissue stiffness map. (A) Schematic depicting 3D tissue stiffness mapping. (B) Three-dimensional rendering of the 20 som. stage pCX-NLS-Cre;mTmG limb bud stiffness map (n=5). (C) H2B-miRFP703 21 som. limb bud three-dimensional mesodermal cell movement trajectories tracked by light sheet live imaging (unit: μm, duration: 3 h). Each dot denotes the last time point of tracking. (D) Schematic describing the definition of cell migration persistence. (E) H2B-miRFP703 21 som. limb bud three-dimensional mesodermal cell migration persistence map overlaid with tissue stiffness map. Grey shade represents the volume mesh of effective stiffness value > 1.5e-3. (F) A representative *z*-section image of 21 som. T-Cre;mTmG. (G) Three-dimensional cell membrane rendering of 21 som. T-Cre;mTmG overlaid with tissue stiffness map. Grey shade represents the volume mesh of effective stiffness value > 1.5e-3. (H) Time-lapse local cell membrane rendering within the three zones defined in F suggesting caged, durotaxis and rearranging behaviours. (I) Proposed zonal cell behaviours as a function of tissue stiffness.

We examined the relation between tissue stiffness and cell movements by performing live, time-lapse light sheet microscopy^2,4^ of embryos harbouring a transgenic far-red nuclear reporter H2B-miRFP703^19^. The resulting movies revealed previously unattainable detail due to our ability to identify and track nearly every nucleus deep within mesoderm with minimal phototoxicity. After correcting for embryo drift by tracking fluorescent beads implanted in the agarose cylinder immediately surrounding the embryo, we observed that large-scale cell movements flowed toward and around the stiff core (Methods, Figure 1C and Movie S2).

To understand the basis of the cell movement pattern, we examined various parameters. The spatial distributions of the mesodermal cell-cell adhesion protein N-cadherin and motor protein phospho-myosin light chain (pMLC) and mean cortical tension as estimated by a transgenic vinculin tension sensor^2^ were spatially uniform within mesoderm (Figure S1E-G). Therefore, the coordinated cell movement pattern we observed is not readily explained by candidate mechanisms such as cell sorting or spatially biased cellular forces. We focused further on the potential role of mesodermal stiffness.

To characterise mesodermal cell movements quantitatively, we calculated cell migration persistence. Persistence is defined as the magnitude of the overall (start-to-finish) cell displacement vector divided by the total length of the displacement trajectory ranging from 0 (random motion) to 1 (straight-line motion)^21^ (Figures 1D and S2A). By registering the tissue stiffness map to the persistence map to examine spatial correlation between the two parameters, we identified a continuum of cell movement types that we categorised into three distinct zones. In the stiff core (Zone 1) where effective stiffness was greater than 1.5e-3 as well as in the soft region furthest from the stiff core (Zone 3), cells exhibited low migration persistence. In contrast, cells immediately surrounding the stiff core where a stiffness gradient was present (Zone 2) migrated with high persistence (Figure 1E). By performing 3D limb bud shape registration, we quantitatively compared the persistence maps measured from different embryos and found the mean standard deviation to be around 10%, indicating that the spatial pattern of cell migration persistence was fairly consistent across different embryos (Methods and Figures S2B-S2D).

To better define the nature of local cell movements in the three zones, we performed live imaging using a transgenic mTmG cell membrane reporter at higher resolution and rendered cells in 4D (Figures 1F-1H and Movies S3-S5). Zonal categorisation was useful for our analysis because it was based on the predominant type of cell movement in each zone. Cells within the stiff core (Zone 1) were least elongated and maintained a stable configuration over time without exchanging neighbours. In Zone 2 where a stiffness gradient dominated, cells exhibited collective movement toward the anteroproximal core. Lastly, in the soft zone (Zone 3), cells displayed abundant protrusive activity and underwent frequent neighbour exchanges. The most basic configuration of rearrangement involved a single cell intercalating into or out of a group of surrounding neighbours that is akin to a 3D T1 exchange we encountered previously in the mandibular arch^2^. In-between zones, we identified intermediate cell behaviours such as durotactic movements encountering caged cells (Movie S6).

Based on these observations, we propose that mesodermal cells transition between 1) caged behaviour in a stiff environment, 2) durotactic movement along a stiffness gradient, and 3) rearrangements in a comparatively soft tissue (Figure 1I).

### Fibronectin is necessary and sufficient for morphogenetic cell movements

Fibronectin is a key component of extracellular matrix (ECM) that determines embryonic tissue stiffness and has been proposed to underlie durotaxis^4,18,22,23^. Unlike other ECM components we previously examined, fibronectin exhibits a spatial distribution that corresponds to mesodermal stiffness gradient^4^. To test whether the relation between tissue stiffness and cell movements is causative *in vivo*, we genetically manipulated fibronectin expression. *Fn*-null mouse embryos develop normally until E8.0 and exhibit severe malformation of mesoderm-derived structures by E8.5^24^. In our study, conditional deletion of homozygous floxed *Fn*^25^ in mesoderm using T-Cre (T-Cre;*Fn*^*f/f*^) resulted in embryos that were progressively smaller compared to their control between E9.5 and E10.5 (Figures S3A and S3B). We confirmed the effective depletion of FN in mutant limb buds by immunostaining (Figures S3C and S3D). Cell proliferation and apoptosis that were examined by pHH3 immunostaining and LysoTracker staining, respectively, were similar between T-Cre;*Fn*^*f/f*^ and wild-type limb buds in ectoderm and mesoderm at 24 som. (∼E9.5) (Figures S3E and S3F). At 34 som. (∼E10.5), a significant reduction in proliferation and an increase in apoptosis were identified in conditional mutants (Figures S3F and S3G).

To examine the effect of fibronectin loss prior to cell proliferation and apoptosis effects, tissue stiffness mapping was performed at 20 som. in embryos that harboured a cell membrane label (T-Cre;*Fn*^*f/f*^;mTmG). Compared to controls, conditional mutants exhibited softer mesoderm that lacked a spatial stiffness bias (Figures 2A and S4A). By live light sheet imaging of T-Cre;*Fn*^*f/f*^;H2B-miRFP703 embryos, the collective pattern of mesodermal cell movements was grossly disrupted with cells in the bulk of the limb bud exhibiting complex, interweaving trajectories (Figure 2B and Movie S7). Correspondingly, cell migration persistence was low throughout the limb bud (Figures 2C and S4B). Time-lapse evaluation of cell membrane-rendered groups of T-Cre;*Fn*^*f/f*^;mTmG cells confirmed wide-spread intercellular rearrangements that we previously observed only in Zone 3 of WT limb buds (Figures 2D, 2E and Movie S8).

**Figure 2.**
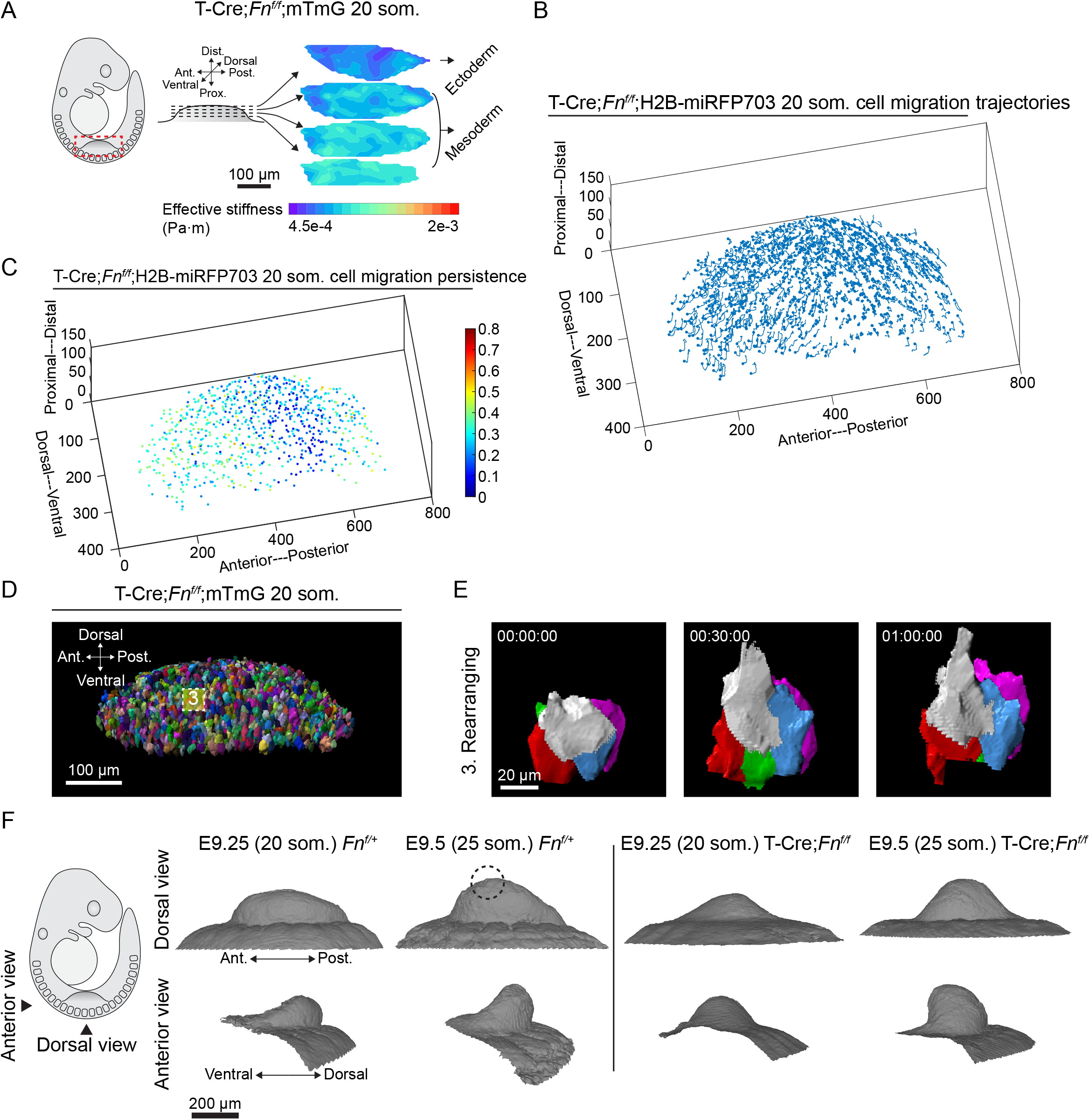
Loss of fibronectin downregulates tissue stiffness and leads to a broadly rearranging state. (A) Three-dimensional rendering of the 20 som. stage T-Cre;*Fn*^*f/f*^;mTmG limb buds stiffness map (n=3). (B) T-Cre;*Fn*^*f/f*^;H2B-miRFP703 20 som. limb bud three-dimensional mesodermal cell movement trajectories tracked by light sheet live imaging (unit: μm, duration: 3 h). Each dot denotes the last time point of tracking. (C) T-Cre;*Fn*^*f/f*^;H2B-miRFP703 20 som. limb bud three-dimensional mesodermal cell migration persistence map. (D) Three-dimensional cell membrane of T-Cre;*Fn*^*f/f*^;mTmG limb bud rendered from light sheet live imaging. (E) Local cell membrane rendering in the dashed square shown in D suggesting a rearranging state. (F) Limb bud shape change from 20 to 25 som. stage *Fn*^*f/+*^ and T-Cre;*Fn*^*f/f*^ embryos reconstructed from optical projection tomography. Dashed circle indicates the location of anteriorly biased peak of the *Fn*^*f/+*^ limb buds.

Using optical projection tomography (OPT), we found that T-Cre;*Fn*^*f/f*^ limb buds developed more symmetrically than those of WT littermates. Mutant limb buds lacked spatial biases such as an anterior peak that normally develops between 20 and 25 som. and a relatively narrow dorsoventral axis (Figures 2F, S4C and S4D). The short AP axis is consistent with previous findings in *Fn*-null embryos^24^ and the wide DV axis resembles that of *Wnt5a* mutants^4,26^. Together, these results indicate that fibronectin is necessary to establish a mesodermal stiffness gradient that regionalises and orients morphogenetic cell movements that shape the limb bud.

To test the effect of excessive fibronectin, we knocked-in a conditional and fluorescently tagged *Fn* transgene to the *Rosa26* locus (R26-CAG-loxP-STOP-loxP-Fn-mScarlet, herein referred to as R26-Fn-mScarlet) using an efficient method of transgenesis 2C-HR-CRISPR^19^ (Methods, Figures 3A and S5A). Overexpression of *Fn* throughout mesoderm using T-Cre abrogated the normally biased FN domain by immunostaining (Figures S5B and S5C) as well as the stiffness gradient by magnetic tweezers measurement, resulting in stiffer tissue throughout the limb bud (Figures 3B and S5D). Live imaging revealed markedly diminished cell movements as indicated by short migration trajectories with low persistence (Figures 3C, 3D, S5E and Movie S9). Accordingly, cells were relatively spherical and in stable configurations that lacked neighbour exchange throughout mesoderm (Figures 3E, 3F and Movie S10). Therefore, excessive fibronectin is sufficient to stiffen tissue in a manner that cages cell movements.

**Figure 3.**
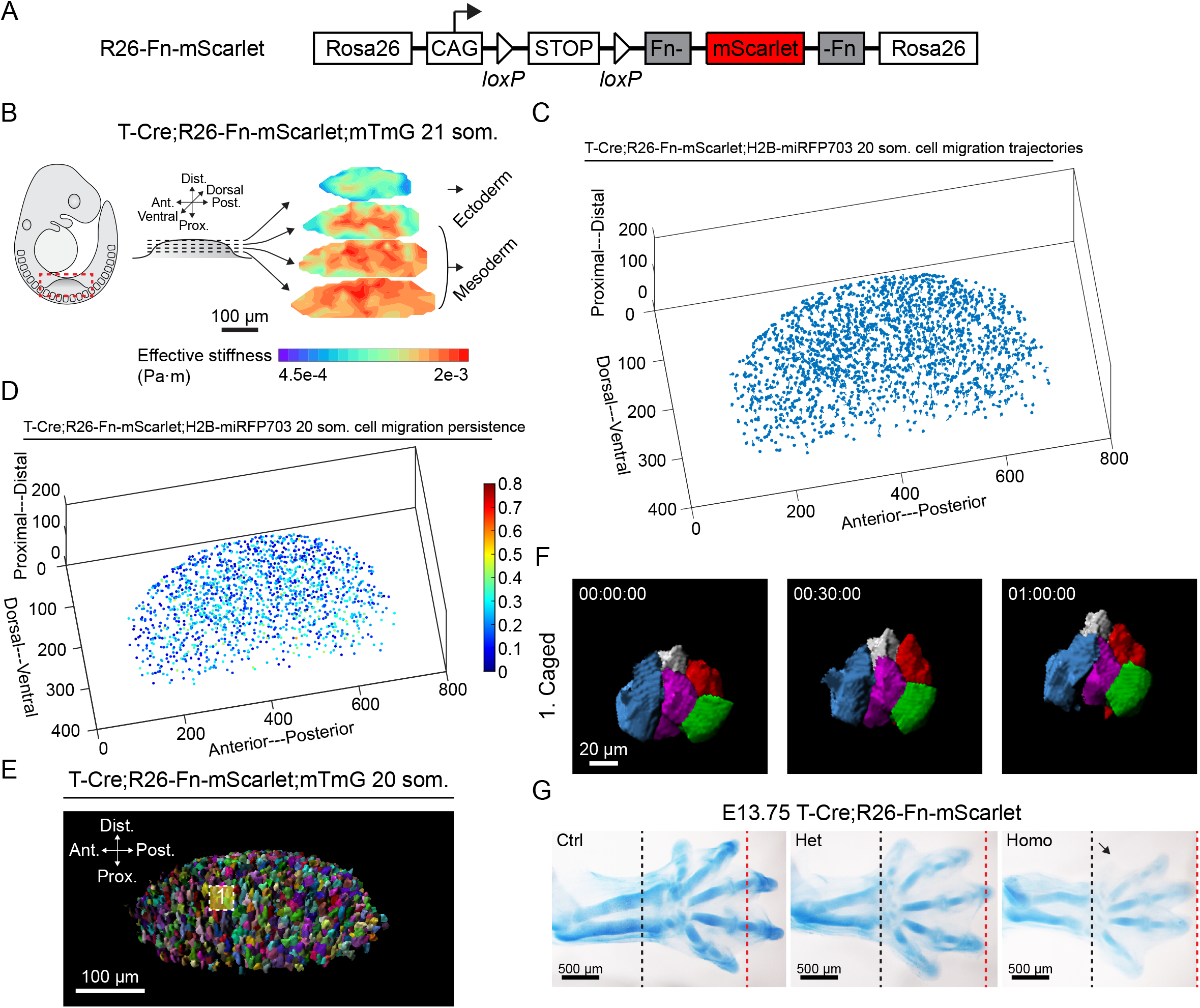
Fibronectin overexpression upregulates tissue stiffness and leads to a broadly caged state. (A) Conditional Rosa26 fibronectin mScarlet knock-in mouse strain. (B) Three-dimensional rendering of the 20 som. stage T-Cre;R26-Fn-mScarlet;mTmG limb buds stiffness map (n=3). (C) T-Cre;R26-Fn-mScarlet;H2B-miRFP703 20 som. limb bud three-dimensional mesodermal cell movement trajectories tracked by light sheet live imaging (unit: μm, duration: 3 h). Each dot denotes the last time point of tracking. (D) T-Cre;R26-Fn-mScarlet;H2B-miRFP703 20 som. limb bud three-dimensional mesodermal cell migration persistence map. (E) Three-dimensional cell membrane of T-Cre;R26-Fn-mScarlet;mTmG limb bud rendered from light sheet live imaging. (F) Local cell membrane rendering suggesting a caged state. (G) E13.75 skeletal staining of mouse forelimb. Black and red dashed lines represent the distal extents of forearm and digits in T-Cre;R26-Fn-mScarlet homo, respectively. Arrow indicates first digit hypoplasia.

Unlike *Fn* mutants, T-Cre;R26-Fn-mScarlet embryos survived until E14.5, permitting assessment of skeletal pattern by cartilage staining. Interestingly, transgenic limbs were disproportionately short with marked radial and first digit hypoplasia (Figure 3G). This dysplasia implies that early morphogenetic movements that generate nuanced tissue shape are fundamentally important for pattern formation. In this case, loss of early anterior tissue bias corresponds to reduction of anterior structures.

### Tissue stiffness predicts the mode of cell movements

The preceding data demonstrate that limb bud mesoderm is a mixed-phase environment. This context is distinct to previously described embryonic tissues such as the zebrafish tail bud and during epiboly in which parameters such as cell packing density^3^ or cell-cell connectivity^12^, respectively, define solid and liquid phases at a transitional critical point. In contrast, mammalian mesoderm exhibits a transitional durotactic zone between solid and liquid phases as a function of local ECM-dependent tissue stiffness. We considered whether rigidity phase transition and durotaxis in this context can be described by a unified mathematical model.

We propose a minimal mathematical description for a stiffness-driven liquid-solid phase transition that includes a description of durotaxis in the intermediate interpolating region within mixed phase mesoderm. Starting with a modified Landau theory for second order phase transitions^27^, we add a term describing the coupling of cellular diffusion to a stiffness gradient (stiffness phase transition model),

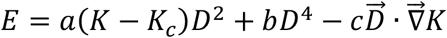

where a, b, and c are positive coefficients. In this equation, the first two terms represent the standard Landau phenomenological description for a phase transition with our postulate that there is a fluid to solid transition driven by tissue stiffness, *K*, above a critical threshold, *Kc*. As in previous works^28^, particle self-diffusivity D (Methods and Figure S6A) is an order parameter for the transition. Diffusivity measures the particle spreading rate in the medium and is an important metric to distinguish the state of matter. It is defined as magnitude of the displacement vector divided by six times the time duration (Methods). The last term in the equation predicts a tilting of the diffusion potential in the direction of the stiffness gradient, resulting in durotaxis (Figure 4A). Our theoretical model suggests that for a zero-stiffness gradient, one would expect the diffusivity to simply transition from a nonzero value at low stiffness (rearranging state) toward zero as stiffness increases (caged state). This postulate is given by a simple sigmoid-type interpolation with the width defining the sharpness of the transition. A real wild-type limb bud, however, does not exhibit uniform stiffness but rather a region of higher stiffness surrounded by a region of lower stiffness with an intermediate region interpolating between the two. It is more akin to a mixed phase situation rather than an equilibrium thermodynamic limit. The interpolating region necessarily has a nonzero stiffness gradient, and so will have some additional dynamics (durotaxis state) and a greater magnitude of diffusivity (Figure 4A).

**Figure 4.**
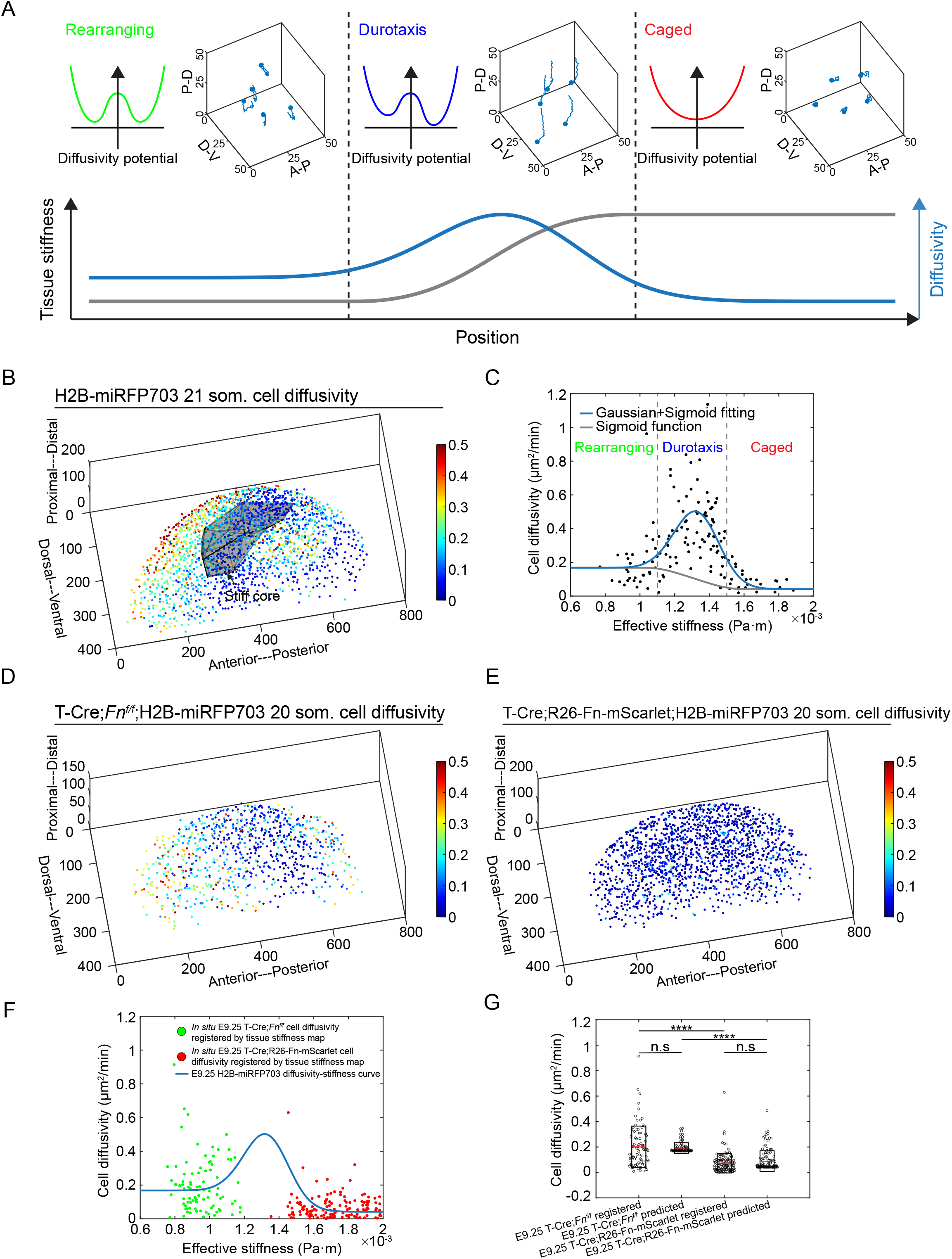
Prediction of cell behaviours from tissue stiffness map. (A) Landau phase diagram as a function of tissue stiffness. (B) H2B-miRFP703 21 som. limb bud three-dimensional mesodermal cell diffusivity map overlaid with tissue stiffness map. Grey shading represents the volume mesh of effective stiffness value > 1.5e-3. (C) Cell diffusivity as a function of tissue stiffness. (D) T-Cre;*Fn*^*f/f*^;H2B-miRFP703 20 som. limb bud three-dimensional mesodermal cell diffusivity map. (E) T-Cre;R26-Fn-mScarlet;H2B-miRFP703 20 som. limb bud three-dimensional mesodermal cell diffusivity map. (F and G) T-Cre;*Fn*^*f/f*^;H2B-miRFP703 and T-Cre;R26-Fn-mScarlet;H2B-miRFP703 limb bud mesodermal cell diffusivity follows the stiffness-phase transition model prediction (two-tailed paired Student’s t-test). n.s, not significant.

An empirical cell diffusivity map of the WT limb bud shows a similar pattern compared to cell migration persistence. The stiff and soft zones both exhibit low diffusivity whereas the stiffness gradient zone has high diffusivity (Figure 4B) in agreement with our stiffness-phase transition model. By registering the WT cell diffusivity map and tissue stiffness map, we obtained the relation between diffusivity and stiffness allowing us to calibrate the stiffness-phase transition model. The gradient of the stiffness roughly follows a Gaussian in the wild-type limb bud (Figures S6B and S6C), so we assumed that diffusivity would likewise diverge from the simple interpolation with a Gaussian line-shape. As such, we fit a sigmoid plus a Gaussian to arrive at our full expected form as shown in Figure 4C. To test the predictive ability of the model, we calculated cell diffusivity maps of *Fn* conditional loss (T-Cre;*Fn*^*f/f*^) and overexpression (T-Cre;R26-Fn-mScarlet embryos) (Figures 4D and 4E) and registered them with tissue stiffness maps. T-Cre;*Fn*^*f/f*^ and T-Cre;R26-Fn-mScarlet embryos both abolished tissue stiffness gradient in limb bud mesoderm lowering cell diffusivity with T-Cre;R26-Fn-mScarlet (i.e., stiffening) being more significant. Therefore, the relationship between diffusivity and stiffness in T-Cre;*Fn*^*f/f*^ and FN overexpressing limb buds follows the prediction of our stiffness-phase transition model (Figures 4F and G). Rigidity phase transition and durotaxis co-exist along a continuum in murine mesoderm.

### Fibronectin is regulated by a feed-forward mechanism involving WNT5A and YAP

Our previous work showed that *Wnt5a* is required for FN expression in the early mouse limb bud^4^. To identify the relevant signalling pathway, we examined pathway components (Figure S7A). The effectors TCF/Lef and the co-activator β-catenin exhibited no spatial-bias in distribution in WT limb buds nor a difference in expression levels between WT and *Wnt5a*^*-/-*^ limb buds (Figures S7B-S7E), suggesting that canonical WNT signalling does not drive FN expression. Similar results were observed for key noncanonical signalling components VANGL2 and c-JUN N-terminal kinase (JNK) as shown in Figures S7F-S7I.

Alternative effectors of WNT5A include YAP/TAZ^29^, although evidence for that pathway *in vivo* is lacking. It has also been shown that *Fn* is a downstream target of YAP *in vitro*^30^. The expression domain of YAP, but not of phospho-YAP that is retained in the cytoplasm ^36^, colocalised with the FN domain (Figures 5A-5C, S8A)). We identified a strong correlation between YAP nuclear/cytoplasmic ratio and FN intensity among individual cells (Methods, Figures 5B and S8B). Next, we examined whether YAP is necessary for FN by performing conditional knockout of floxed *Yap/Taz* using T-Cre. In *Yap/Taz* mutant limb buds, FN immunostain intensity was broadly diminished (Figures 5D and 5E) and limb bud shapes resembled those of *Fn* conditional knockouts with greater severity at 20 and 25 som. stages (Figure 5F). These data suggest YAP is an upstream regulator of FN *in vivo*.

**Figure 5.**
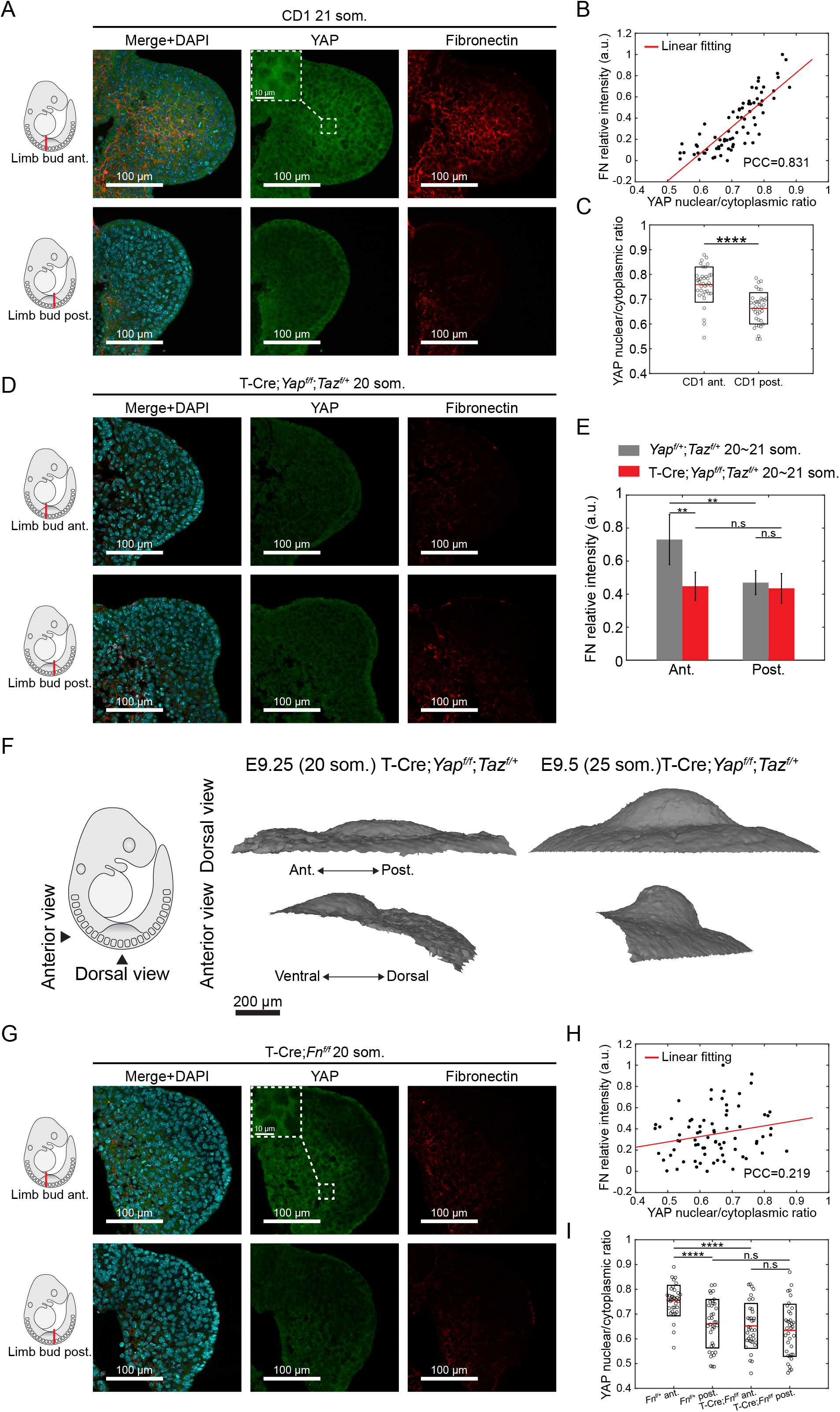
YAP regulates fibronectin expression via a feedback mechanism. (A) Transverse sections of 21 som. CD1 embryos at forelimb anterior and posterior regions. Sections were stained with DAPI (cyan) anti-YAP (green) and anti-fibronectin antibody (red). (B) Relative fibronectin fluorescence intensity vs. YAP nuclear/cytoplasmic ratio of 20∼21 som. CD1 embryos (n=3) at forelimb anterior region. PCC: Pearson correlation coefficient. (C) YAP nuclear/cytoplasmic ratio of 20∼21 som. CD1 embryos (n=3) at forelimb anterior and posterior regions (two-tailed unpaired Student’s t-test, \*\*\*\**P* < 0.0001). (D) Transverse sections of 20 som. T-Cre;*Yap*^*f/f*^;*Taz*^*f/+*^ embryos at forelimb anterior and posterior regions. Sections were stained with DAPI (cyan) anti-YAP (green) and anti-fibronectin antibody (red). (E) Relative fibronectin fluorescence intensity in 20∼21 som. *Yap*^*f/+*^;*Taz*^*f/+*^ (n=5) vs. T-Cre;*Yap*^*f/f*^;*Taz*^*f/+*^ (n=3) embryos at forelimb anterior and posterior regions (two-tailed unpaired Student’s t-test, \*\**P* < 0.01). (F) Limb bud shape change from 20 to 25 som. stage T-Cre;*Yap*^*f/f*^;*Taz*^*f/+*^ embryos reconstructed from optical projection tomography. (G) Transverse sections of 20 som. T-Cre;*Fn*^*f/f*^ embryos at forelimb anterior and posterior regions. Sections were stained with DAPI (cyan) anti-YAP (green) and anti-fibronectin antibody (red). (H) Relative fibronectin fluorescence intensity vs. YAP nuclear/cytoplasmic ratio in 20∼21 som. T-Cre;*Fn*^*f/f*^ embryos (n=3) at forelimb anterior region. (I) YAP nuclear/cytoplasmic ratio of 20∼21 som. *Fn*^*f/+*^ (n=3) vs. T-Cre;*Fn*^*f/f*^ (n=3) embryos at forelimb anterior and posterior regions (two-tailed unpaired Student’s t-test, \*\*\*\**P* < 0.0001). n.s, not significant. Error bars indicate s.d..

Various *in vitro* studies have shown that YAP is translocated to the nucleus on a stiff substrate^31–33^. We found that, in T-Cre;*Fn*^*f/f*^ limb buds, YAP nuclear/cytoplasmic ratio was diminished and its colocalisation with fibronectin was disrupted (Figures 5G-5I). In *Wnt5a* mutants, both YAP and FN were diminished (Figures 6A-6C). The *Wnt5a* expression domain and level of expression, however, were unaffected in *Yap*-CKO limb buds, indicating an upstream role of *Wnt5a* for YAP activation (Figures 6D and 6E). Interestingly, in T-Cre;*Fn*^*f/f*^ limb buds *Wnt5a* expression level, but not its expression domain, was downregulated by quantitative RT-PCR (Figure 6F). Therefore, FN feeds back to promote nuclear translocation of YAP and expression of *Wnt5a* (Figure 6G).

**Figure 6.**
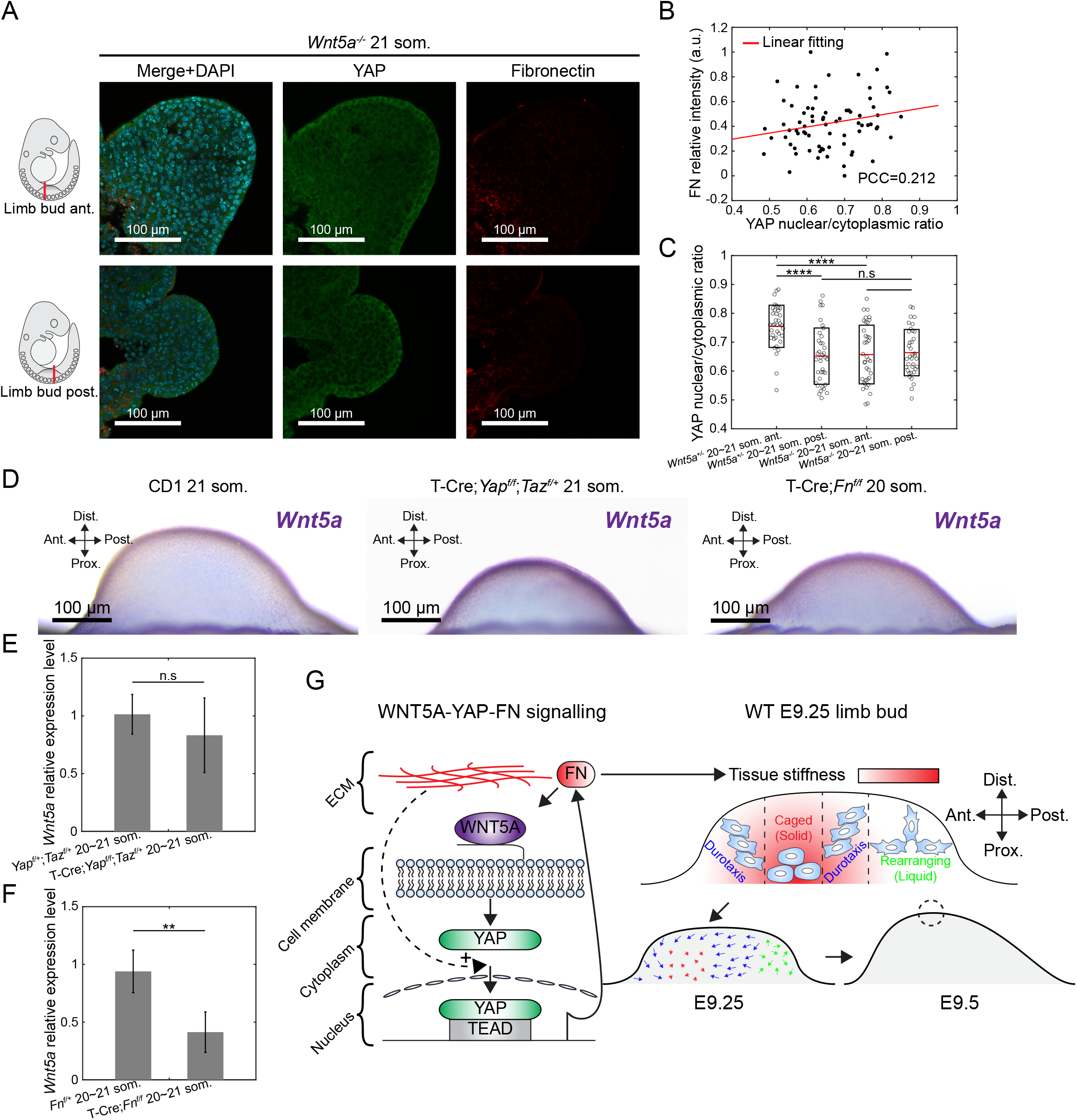
Fibronectin scaffold stiffness orchestrates distinct cell movements via a feedback mechanism involving *Wnt5a*-YAP. (A) Transverse sections of 21 som. *Wnt5a*^-/-^ embryos at forelimb anterior and posterior regions. Sections were stained with DAPI (cyan) anti-YAP (green) and anti-fibronectin antibody (red). (B) Relative fibronectin fluorescence intensity vs. YAP nuclear/cytoplasmic ratio of 20∼21 som. *Wnt5a*^-/-^ embryos (n=3) at forelimb anterior region. PCC: Pearson correlation coefficient. (C) YAP nuclear/cytoplasmic ratio of 20∼21 som. *Wnt5a*^+/-^ (n=3) vs. *Wnt5a*^-/-^ (n=3) embryos at forelimb anterior and posterior regions (two-tailed unpaired Student’s t-test, \*\*\*\**P* < 0.0001). (D) Coronal views of *Wnt5a* expression domain in E9.25 CD1, T-Cre;*Yap*^*f/f*^;*Taz*^*f/+*^ and T-Cre;*Fn*^*f/f*^ forelimbs. (E and F) Wnt5a mRNA level in T-Cre;*Yap*^*f/f*^;*Taz*^*f/+*^ and T-Cre;*Fn*^*f/f*^ forelimbs comparing to *Yap*^*f/+*^;*Taz*^*f/+*^ and *Fn*^*f/+*^, respectively (two-tailed unpaired Student’s t-test, \*\**P* < 0.01). n.s, not significant. Error bars indicate s.d.. (G) Schematic model representing the WNT5A-YAP-FN feedback signalling pathway establishes the tissue stiffness that orchestrates cell movements to drive limb bud shape change and affect cartilage formation.

## DISCUSSION

Our data show that control of morphogenetic cell movements is achieved by tuning fibronectin-dependent tissue stiffness. They imply the location of a stiffness gradient organises cell movements, tissue shape and subsequent skeletal pattern. In contrast to the proximally stiff, distally soft limb bud, the murine mandibular arch exhibits distally-biased stiffness where cells are caged and proximal soft mesenchyme where cells intercalate^2,23^. Stiff regions promote structural stability whereas soft regions permit deformability and elongation, resulting in different 3D mesodermal tissue shapes depending on stiffness geometry.

Our stiffness-phase transition model unifies two concepts that were previously considered separately. Durotaxis takes place in an interpolating region in mixed phase tissue. It will be interesting to determine whether transitions in cell behaviour along this continuum occur at stiffness thresholds that are absolute or relative. Examination of tissues or embryos that are substantially larger or smaller than the limb bud with different tissue properties will be instructive in this regard and potentially useful for tissue engineering applications.

We used the term durotaxis to explain our results because that term accurately reflects our stiffness measurements. An alternative interpretation of our fibronectin data is that cells moved up a gradient of integrin-fibronectin binding sites (haptotaxis). Unlike *in vitro* environments in which substrate stiffness and binding site density can be modified separately by altering polymer concentration or ECM protein coating, ECM-dependent stiffness and binding site density are not readily distinguishable *in vivo*. Previous inactivation of a fibronectin binding motif (RGD of FN-III) for α51/αϖ integrins did not compromise FN matrix assembly^34^ and resulted in a less severe phenotype than Fn null embryos^24^. However, there were marked mesodermal anomalies including hypoplasia of the limb bud, raising the possibility that cells can’t migrate effectively in the absence of ECM binding sites despite the presence of a matrix. We infer that durotaxis and haptotaxis are inherently coupled or interdependent *in vivo*.

Previous studies have shown that both WNT ligands and YAP are capable of driving fibronectin expression^35,36^. In limb bud mesoderm YAP acts as a downstream effector of *Wnt5a* to regulate fibronectin expression which in turn promotes nuclear translocation of YAP and *Wnt5a* expression. Fibronectin feedback can be transcriptional, post-transcriptional or combined. Since nuclear shuttling of YAP has long been shown to respond to changes in environmental stiffness *in vitro*, mechanotransduction represents an obvious candidate mechanism^31–33^. Dynamic changes in YAP location during development were recently confirmed using an endogenous YAP reporter^37^. Interplay between biochemical and mechanical signals likely controls a wide range of developmental events.

Our work identifies a fundamental layer of morphogenetic regulation by revealing that durotaxis interpolates within mixed phase tissue. This principle may be widely applicable but regulated by different ECM components or control variables in other contexts. Identifying key control variables that underlie phase behaviour can potentially advance our morphogenetic understanding of congenital malformations and tissue engineering strategies in which form is critically important for function.

## MATERIALS AND METHODS

### Mouse strains

Analysis was performed using the following mouse strains: pCX-NLS-Cre^38^, mTmG^39^ [The Jackson Laboratory: Gt(ROSA)26Sor^tm4(ACTB-tdTomato-EGFP)Luo^/J)], R26-CAG-H2B-miRFP703^19^ [The Jackson Laboratory: Gt(ROSA)26Sor^em1.1(CAG-RFP*)Jrt^/J], T-Cre^40^, Fn1 flox^25^ [The Jackson Laboratory: B6;129-Fn1^tm1Ref^/J], Rosa26-CAG-LSL-Fn-mScarlet (R26-Fn-mScarlet)-see below, Yap flox;Taz flox^41^ [The Jackson Laboratory: Wwtr^1tm1Hmc^ Yap1^tm1Hmc^/WranJ], Wnt5a^+/−42^ and TCF/Lef:H2B-GFP^43^. To generate conditional mutant embryos, flox/flox females carrying the appropriate fluorescent reporter were bred to flox/+; Cre males. All strains were outbred to CD1 except for Fn1 flox, which is C57BL/6J background. All animal experiments were performed in accordance with protocols approved by the Animal Care Committee of the Hospital for Sick Children Research Institute.

### Generation of Rosa26-CAG-LSL-Fn-mScarlet mouse line

The Fn-mScarlet transgene was designed according to a previous publication demonstrating normal expression and secretion of Fn-GFP fusion protein^44^. The coding sequence of a bright red fluorescence protein mScarlet was placed between the FN-III domains 3 and 4 within the full-length cDNA sequence of the longest isoform of mouse Fn1. The full length Fn-mScarlet cDNA was synthesized by Epoch Inc and then inserted into the MluI restriction site of the pR26 CAG AsisL/MluI plasmid (Addgene 74268, kind gift from Ralph Kuehn)^45^. The mouseline was generated by 2C-HR-CRISPR following our published protocol^19,46,47^. A positive founder was outcrossed for 3 generations to remove any potential off-target modification from CRISPR genome editing before breeding to establish desired genotypes. The Rosa26-CAG-LSL-Fn-mScarlet are homozygous viable, healthy, and fertile. The mouse line is maintained by breeding homozygous mice.

### 3D Tissue stiffness mapping

3D tissue stiffness mapping was conducted using a 3D magnetic device that we developed previously. Briefly, functionalised magnetic beads (Dynabead M280) were microinjected into the limb bud field using a microinjector (CellTram 4r Oil; Eppendor) at anterior, middle, and posterior regions. Multiple depositions were made during one penetration and withdrawal of the needle to distribute the magnetic beads at different depths within the limb bud. The embryo with magnetic beads injected was placed into the customized imaging chamber and immobilised by DMEM without phenol red containing 12.5% rat serum and 1% low-melt agarose (Invitrogen). The temperature of the imaging chamber was maintained at 37 °C with 5% CO_2_. Prior to stiffness mapping, a Z scan was taken to record the bead locations within the limb bud. Magnetic beads were then actuated by the magnetic device and bead displacements were recorded by spinning-disk confocal microscopy at the highest frame rate. The bead displacement (in the direction of the magnetic force) was tracked with subpixel resolution and fitted using Zener model with a serial dashpot in MATLAB to extract the stiffness value. The tissue stiffness mapping dataset was rendered in 3D using a customised python program.

### 3D Limb bud registration

Our 3D limb bud registration was inspired by previous work on segmentation of the full lower limb in computed tomography (CT) and construction of an average embryo from multiple reconstructed embryos. The 3D point cloud of mouse limb bud to be registered (source) was first aligned with the reference (target) in AP and DV axes by translation and rotation. The centroid of the source was then merged with the target. An iterative closest point (ICP) based non-rigid registration algorithm was applied until reaching a shape difference below 5%. The shape difference is defined as

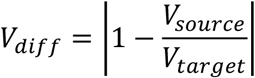

where *V*_*source*_ and *V*_*target*_ are the volumes of source and target, respectively.

### Live, time-lapse light-sheet microscopy

Three-dimensional time-lapse microscopy was performed on a Zeiss Lightsheet Z.1 microscope. Embryos were suspended in a solution of DMEM without phenol red containing 12.5% filtered rat serum, 1% low-melt agarose (Invitrogen), and 2% fluorescent beads (1:500, diameter: 500 nm; Sigma-Aldrich) that were used for drift-compensation in a glass capillary tube. Once the agarose solidified, the capillary was submerged into an imaging chamber containing DMEM without phenol red, and the agarose plug was partially extruded from the glass capillary tube until the portion containing the embryo was completely outside of the capillary. The temperature of the imaging chamber was maintained at 37 °C with 5% CO2. Images were acquired using a 20×/0.7 objective with dual-side illumination, and a z-interval of 0.5μm. Images were acquired for 3 to 4 h with 5 min intervals.

### Cell migration tracking and analysis

The light-sheet time-lapse dataset was rendered in Imaris (Bitplane). The positions of cell nuclei and fluorescent beads were tracked over time using an autoregressive motion algorithm. Ectodermal and mesodermal cells were separated based on mean thresholding of fluorescence intensity. Cells undergoing division (based on the morphology of the cell nuclei) were excluded from cell migration tracking. The tracking dataset was then imported into MATLAB (MathWorks), and drift compensation was performed by translation and rotation based on the displacement of fluorescent beads. Cell migration metrics were calculated using the drift-compensated dataset. Cell migration persistence *P* was calculated by dividing vector length by the length of the trajectory

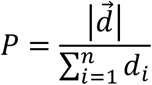

As in previous works^28^, we defined particle self diffusivity D as

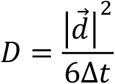

where *t* is the time in mins.

To evaluate the cross-embryo difference of the calculated metrics, multiple limb bud datasets were registered to the reference. For each cell in the reference dataset, the closest cell was searched in other limb bud datasets within the distance of 20 μm. The standard deviation of each metric was calculated among the cells from different embryos at the proximity of reference location.

### Fluorescence lifetime imaging microscopy

Fluorescence lifetime microscopy (FLIM) was performed on a Nikon A1R Si laser scanning confocal microscope equipped with PicoHarp 300 TCSPC module and a 440 nm pulsed diode laser (Picoquant). Data were acquired with a 40×/1.25 water immersion objective with a pixel dwell time of 12.1 µs/pixel, 512×512 resolution, and a repetition rate of 20 MHz. Fluorescence emission of mTFP1 was collected through a 482/35 bandpass filter. Embryos were prepared in 50/50 rat serum/DMEM and imaged in a humidified chamber at 37 °C in 5% CO_2_. Lifetime analysis was performed using FLIMvivo^48^.

### Optical projection tomography

Mouse embryos were harvested and fixed in 4% paraformaldehyde overnight at 4 °C. Optical projection tomography was performed using a system that was custom-built and is fully described elsewhere^49^. Three-dimensional datasets were reconstructed from auto-fluorescence projection images acquired during a 25-min scan period at an isotropic voxel size of 4.5μm. The limb bud structure was segmented from the embryo and rendered in MeshLab.

### Immunofluorescence

Cryosection immunofluorescent staining of embryos was performed as previously described^4^. Dissected mouse embryos were fixed overnight in 4% paraformaldehyde in PBS followed by three washes for 5 min in PBS. Fixed embryos were embedded in 7.5% gelatin/15% sucrose and sectioned into 10-μm slices using a Leica CM1800 cryostat. Sections were washed twice for 5 min. in Milli-Q and once for 5 min in PBS, permeabilised in 0.1% Triton X-100 in PBS for 20 min, and blocked in 5% normal donkey serum (in 0.05% Triton X-100 in PBS) for 1 h. Sections were incubated in primary antibody overnight at 4 °C followed by four 10-min washes in 0.05% Triton X-100 in PBS, then incubated in secondary antibody (1:1,000) for 1 h at room temperature. Finally, sections were washed three times for 5 min in 0.05% Triton X-100 in PBS and twice for 5 min in PBS. Images were acquired using a Nikon A1R Si Point Scanning Confocal microscope at 20× or 40× magnification, and analysis was performed using ImageJ.

### Whole-mount immunofluorescence

Whole-mount immunofluorescent staining of embryos was performed as previously described^2^. Briefly, mouse embryos were collected and fixed overnight in 4% PFA in PBS followed by 3 washes in PBS. Embryos were permeabilised in 0.1% Triton X-100 in PBS for 20 min and blocked in 5% normal donkey serum (in 0.05% Triton X-100 in PBS) for 1 h. Embryos were incubated in primary antibody for 5 h at room temperature, followed by overnight incubation at 4 °C. Embryos were washed 4×20 min with 0.05% Triton X-100 in PBS and incubated in secondary antibody for 3-5 h at room temperature.

Embryos were washed 4×20 min, followed by a final wash overnight at 4 °C, and stored in PBS. Images were acquired using a Nikon A1R Si Point Scanning Confocal microscope at 20× or 40× magnification, and analysis was performed using Imaris (Bitplane).

### Apoptosis detection

LysoTracker Red DND-99 (ThermoFisher Lot L7528) was diluted to 2μM in DMEM containing 50% rat serum. Embryos were placed in media and incubated in a roller culture apparatus for 1 h. The temperature was maintained at 37 °C with 5% CO2. Embryos were washed three times with PBS after staining to remove LysoTracker surplus then fixed overnight in 4% paraformaldehyde in PBS followed by three washes for 5 min. in PBS. Images were acquired using a Nikon A1R Si Point Scanning Confocal microscope at 20X magnification, and analysis was performed using ImageJ.

### Whole-mount skeleton staining

Dissected E13.5 mouse embryos in glass scintillation vials were fixed overnight in 70% ethanol at 4 °C. The 70% ethanol was removed and replaced with 95% ethanol for 1 h at room temperature. The embryos were then permeabilized by acetone for 1 h followed by Alcian blue colorimetric reaction for 1 h at room temperature. The embryos were kept in 1% KOH overnight at 4 °C for clearing. Finally, embryos were transferred to 50 % glycerol:50 % (1 %) KOH solution for 2 h at room temperature and kept in 100 % glycerol for long term storage. For imaging, embryos were transferred to a solution containing 45% sucrose and 40% urea for better imaging quality. The images were captured using Leica stereo microscope MSV269.

### Whole-mount *in-situ* hybridisation

Whole mount *in situ* hybridisation was performed after fixation in 4% paraformaldehyde followed by dehydration through a methanol series. Embryos were bleached then underwent digestion in proteinase K, hybridization with probe in formamide, SSC, SDS and heparin, treatment with anti-Digoxygenin-AP, and colorimetric reaction. Wildtype and mutant littermate embryos were processed identically in the same assay for comparison. The images were captured using Leica stereo-microscope MSV269.

### Antibodies

Primary antibodies included fibronectin (1:100; Abcam Lot ab2413), pHH3 (1:250; Cell Signaling Lot 9706), N-cadherin (1:200; BD Biosciences Lot 610920), beta-catenin (1:200; Abcam Lot ab59430), Vangl2 (1:200; Sigma Lot HPA027043), p-SAPK/JNK (1:200; Cell Signaling Lot 9255), YAP (1:200; Abcam Lot ab56701) and phosphor-YAP (1:200; Cell Signaling Lot 4911). Secondary antibodies were purchased from Abcam and used at 1:1,000 dilutions: Goat Anti Mouse IgG AF 488 (ab150113), Goat Anti Rabbit IgG AF 488 (ab150077), Goat Anti Rabbit IgG AF 568 (ab175471).

### Primers

Primers used in this study included R26 Fn-mScarlet gt Fwd: GTACAACATCACTATCTATGCTGTGGA R26 Fn-mScarlet gt Rev: CGTCCTCGAAGTTCATCACGC R26 WT Fwd: CGTGCAAGTTGAGTCCATCCGCC R26 WT Rev: ACTCCGAGGCGGATCACAAGCA Wnt5a Fwd: CTCGGGTGGCGACTTCCTCTCCG Wnt5a Rev: CTATAACAACCTGGGCGAAGGAG

## Data Availability

The data that support the findings of this study are available at https://figshare.com/articles/dataset/Durotaxis_bridges_phase_transition_as_a_function_of_tissue_stiffness_in_vivo.

## Code Availability

All custom codes used in this paper are available at https://github.com/MinZhuUOTSickKids/ Durotaxis-bridges-phase-transition-as-a-function-of-tissue-stiffness-in-vivo.

## ACKNOWLEDGEMENTS

This work was funded by the Canadian Institutes of Health Research (168992) to SH/YS the Canada First Research Excellence Fund/Medicine by Design (MbDGQ-2021-04) to SH/YS/SG.

## AUTHOR CONTRIBUTIONS

M.Z. designed and performed most experiments, modelling, and wrote the manuscript; B.G. generated Fn overexpression strain; E.T. performed mathematical modelling; H.T. and T.Y. technically supported the murine experiments; K.Z. technically supported tissue stiffness measurements; J.R. supervised generation of the Fn overexpression strain; Y.S. and S.H. co-conceived and co-funded the project and edited the manuscript.

## DECLARATION OF INTERESTS

The authors declare no competing interests.

## SUPPLEMENTARY MOVIE LEGENDS

**Movie S1**

Three-dimensional limb bud shape registration using a customised MATLAB program.

**Movie S2**

Three-dimensional rendering and time lapse movie of a 21 somite H2B-miRFP703 (WT) transgenic embryonic limb bud imaged by light sheet microscopy. The view is dorsal, anterior is to the left, and distal is upwards. Any movement of the whole embryo within the imaging chamber is not confounding here as fluorescent beads were embedded in agarose immediately surrounding the embryo to allow for drift compensation. In this whole limb bud view, cells within the stiff anteroproximal core are relatively static whereas surrounding cells move toward that core.

**Movie S3**

Local cell membrane rendering and time lapse movie of a 21 somite T-Cre;mTmG (WT) transgenic embryonic limb bud within the stiff core. Although individual cell membranes fluctuate to some degree, cell neighbour relationships remain fixed, reflecting a caged state.

**Movie S4**

Local cell membrane rendering and time lapse movie of a 21 somite T-Cre;mTmG (WT) transgenic embryonic limb bud within the stiffness gradient between stiff and soft regions. The elongated nature and coordinated movement of cells are distinct within this zone. Note that drift compensation has been performed and displacements reflect actual cell movements within mesoderm although the background has been blacked out to permit 3D visualisation of cell membranes.

**Movie S5**

Local cell membrane rendering and time lapse movie of a 21 somite T-Cre;mTmG (WT) transgenic embryonic limb bud in the soft zone. Cell shape fluctuations are robust and the blue cell intercalates down into a group of surrounding cells, a process that reflects a 3D version of a T1 intercalation.

**Movie S6**

Local cell membrane rendering and time lapse movie of a 21 somite T-Cre;mTmG (WT) transgenic embryonic limb bud at the junction of the stiff core and the stiffness gradient. The white cell exhibits directional movement that halts as it encounters caged cells.

**Movie S7**

Three-dimensional rendering and time lapse movie of a 20 somite T-Cre;*Fn*^*f/f*^;H2B-miRFP703 transgenic embryonic limb bud imaged by light sheet microscopy. The view is dorsal, anterior is to the left, and distal is upwards. In the absence of FN, cells move globally but lack coordination.

**Movie S8**

Local cell membrane rendering and time lapse movie of a 20 somite T-Cre;*Fn*^*f/f*^;mTmG transgenic embryonic limb bud within the normally stiff zone. In the absence of FN, cells that normally are caged exhibit robust shape fluctuations and the green cell intercalates then withdraws from between the red, white and blue cells. This type of intercellular movement is normally seen only in the soft region.

**Movie S9**

Three-dimensional rendering and time lapse movie of a 20 somite T-Cre;R26-Fn-mScarlet;H2B-miRFP703 transgenic embryonic limb bud imaged by light sheet microscopy. The view is dorsal, anterior is to the left, and distal is upwards. In the presence of excessive FN, cells are globally relatively static and uncoordinated.

**Movie S10**

Local cell membrane rendering and time lapse movie of 20 somite T-Cre;R26-Fn-mScarlet;mTmG transgenic embryonic limb bud within the normally soft zone. In the presence of excessive FN, cell intercalations or directional movements are absent.

## Notes

### Competing Interest Statement

The authors have declared no competing interest.

## REFERENCES

1. Boehm, B., Westerberg, H., Lesnicar-Pucko, G., Raja, S., Rautschka, M., Cotterell, J., Swoger, J., and Sharpe, J. (2010). The role of spatially controlled cell proliferation in limb bud morphogenesis. PLoS Biol. 10.1371/journal.pbio.1000420.

2. Tao, H., Zhu, M., Lau, K., Whitley, O.K.W., Samani, M., Xiao, X., Chen, X.X., Hahn, N.A., Liu, W., Valencia, M., et al. (2019). Oscillatory cortical forces promote three dimensional cell intercalations that shape the murine mandibular arch. Nat Commun. 10.1038/s41467-019-09540-z.

3. Mongera, A., Rowghanian, P., Gustafson, H.J., Shelton, E., Kealhofer, D.A., Carn, E.K., Serwane, F., Lucio, A.A., Giammona, J., and Campàs, O. (2018). A fluid-to-solid jamming transition underlies vertebrate body axis elongation. Nature. 10.1038/s41586-018-0479-2.

4. Zhu, M., Tao, H., Samani, M., Luo, M., Wang, X., Hopyan, S., and Sun, Y. (2020). Spatial mapping of tissue properties in vivo reveals a 3D stiffness gradient in the mouse limb bud. Proc Natl Acad Sci U S A. 10.1073/pnas.1912656117.

5. Cetera, M., Leybova, L., Woo, F.W., Deans, M., and Devenport, D. (2017). Planar cell polarity-dependent and independent functions in the emergence of tissue-scale hair follicle patterns. Dev Biol 428, 188–203. 10.1016/J.YDBIO.2017.06.003.

6. Scott, I.C. (2012). Life Before Nkx2.5: Cardiovascular Progenitor Cells: Embryonic Origins and Development. Curr Top Dev Biol 100, 1–31. 10.1016/B978-0-12-387786-4.00001-4.

7. Barriga, E.H., and Theveneau, E. (2020). In vivo Neural Crest Cell Migration Is Controlled by “Mixotaxis.” Front Physiol 11. 10.3389/fphys.2020.586432.

8. Minegishi, K., Hashimoto, M., Ajima, R., Takaoka, K., Shinohara, K., Ikawa, Y., Nishimura, H., McMahon, A.P., Willert, K., Okada, Y., et al. (2017). A Wnt5 Activity Asymmetry and Intercellular Signaling via PCP Proteins Polarize Node Cells for Left-Right Symmetry Breaking. Dev Cell. 10.1016/j.devcel.2017.02.010.

9. Xiong, F., Tentner, A.R., Huang, P., Gelas, A., Mosaliganti, K.R., Souhait, L., Rannou, N., Swinburne, I.A., Obholzer, N.D., Cowgill, P.D., et al. (2013). Specified neural progenitors sort to form sharp domains after noisy Shh signaling. Cell 153. 10.1016/j.cell.2013.03.023.

10. Sunyer, R., Conte, V., Escribano, J., Elosegui-Artola, A., Labernadie, A., Valon, L., Navajas, D., García-Aznar, J.M., Muñoz, J.J., Roca-Cusachs, P., et al. (2016). Collective cell durotaxis emerges from long-range intercellular force transmission. Science (1979) 353, 1157–1161. 10.1126/science.aaf7119.

11. Barriga, E.H., Franze, K., Charras, G., and Mayor, R. (2018). Tissue stiffening coordinates morphogenesis by triggering collective cell migration in vivo. Nature 554, 523–527. 10.1038/nature25742.

12. Petridou, N.I., Corominas-Murtra, B., Heisenberg, C.P., and Hannezo, E. (2021). Rigidity percolation uncovers a structural basis for embryonic tissue phase transitions. Cell 184, 1914-1928.e19. 10.1016/J.CELL.2021.02.017.

13. Lenne, P.F., and Trivedi, V. (2022). Sculpting tissues by phase transitions. Nature Communications 2022 13:1 13, 1–14. 10.1038/s41467-022-28151-9.

14. Merkel, M., and Manning, M.L. (2018). A geometrically controlled rigidity transition in a model for confluent 3D tissues. New J Phys. 10.1088/1367-2630/aaaa13.

15. Kim, S., Pochitaloff, M., Stooke-Vaughan, G.A., and Campàs, O. (2021). Embryonic tissues as active foams. Nature Physics 2021 17:7 17, 859–866. 10.1038/s41567-021-01215-1.

16. Parada, C., Banavar, S.P., Khalilian, P., Rigaud, S., Michaut, A., Liu, Y., Joshy, D.M., Campàs, O., and Gros, J. (2022). Mechanical feedback defines organizing centers to drive digit emergence. Dev Cell 57, 854-866.e6. 10.1016/J.DEVCEL.2022.03.004.

17. Sunyer, R., and Trepat, X. (2020). Durotaxis. Current Biology. 10.1016/j.cub.2020.03.051.

18. Shellard, A., and Mayor, R. (2021). Collective durotaxis along a self-generated stiffness gradient in vivo. Nature 600. 10.1038/s41586-021-04210-x.

19. Gu, B., Posfai, E., and Rossant, J. (2018). Efficient generation of targeted large insertions by microinjection into two-cell-stage mouse embryos. Nat Biotechnol. 10.1038/nbt.4166.

20. Audenaert, E.A., van Houcke, J., Almeida, D.F., Paelinck, L., Peiffer, M., Steenackers, G., and Vandermeulen, D. (2019). Cascaded statistical shape model based segmentation of the full lower limb in CT. Comput Methods Biomech Biomed Engin 22. 10.1080/10255842.2019.1577828.

21. Gorelik, R., and Gautreau, A. (2014). Quantitative and unbiased analysis of directional persistence in cell migration. Nat Protoc 9. 10.1038/nprot.2014.131.

22. Hartman, C.D., Isenberg, B.C., Chua, S.G., and Wong, J.Y. (2016). Vascular smooth muscle cell durotaxis depends on extracellular matrix composition. Proc Natl Acad Sci U S A 113, 11190–11195. 10.1073/pnas.1611324113.

23. Zhu, M., Zhang, K., Tao, H., Hopyan, S., and Sun, Y. (2020). Magnetic Micromanipulation for In Vivo Measurement of Stiffness Heterogeneity and Anisotropy in the Mouse Mandibular Arch. Research. 10.34133/2020/7914074.

24. George, E.L., Georges-Labouesse, E.N., Patel-King, R.S., Hynes, R.O., and Rayburn, H. (1993). Defects in mesoderm, neural tube and vascular development in mouse embryos lacking fibronectin. Development.

25. Sakai, T., Johnson, K.J., Murozono, M., Sakai, K., Magnuson, M.A., Wieloch, T., Cronberg, T., Isshiki, A., Erickson, H.P., and Fässler, R. (2001). Plasma fibronectin supports neuronal survival and reduces brain injury following transient focal cerebral ischemia but is not essential for skin-wound healing and hemostasis. Nat Med. 10.1038/85471.

26. Lau, K., Tao, H., Liu, H., Wen, J., Sturgeon, K., Sorfazlian, N., Lazic, S., Burrows, J.T.A., Wong, M.D., Li, D., et al. (2015). Anisotropic stress orients remodelling of mammalian limb bud ectoderm. Nat Cell Biol 17, 569–579. 10.1038/ncb3156.

27. Landau, L.D., and Lifshitz, E.M. (2013). Statistical Physics: Volume 5 (Elsevier).

28. Bi, D., Yang, X., Marchetti, M.C., and Manning, M.L. (2016). Motility-driven glass and jamming transitions in biological tissues. Phys Rev X. 10.1103/PhysRevX.6.021011.

29. Park, H.W., Kim, Y.C., Yu, B., Moroishi, T., Mo, J.S., Plouffe, S.W., Meng, Z., Lin, K.C., Yu, F.X., Alexander, C.M., et al. (2015). Alternative Wnt Signaling Activates YAP/TAZ. Cell 162. 10.1016/j.cell.2015.07.013.

30. Nardone, G., Oliver-De La Cruz, J., Vrbsky, J., Martini, C., Pribyl, J., Skládal, P., Pešl, M., Caluori, G., Pagliari, S., Martino, F., et al. (2017). YAP regulates cell mechanics by controlling focal adhesion assembly. Nat Commun 8. 10.1038/ncomms15321.

31. Dupont, S., Morsut, L., Aragona, M., Enzo, E., Giulitti, S., Cordenonsi, M., Zanconato, F., le Digabel, J., Forcato, M., Bicciato, S., et al. (2011). Role of YAP/TAZ in mechanotransduction. Nature 474. 10.1038/nature10137.

32. Meli, V.S., Atcha, H., Veerasubramanian, P.K., Nagalla, R.R., Luu, T.U., Chen, E.Y., Guerrero-Juarez, C.F., Yamaga, K., Pandori, W., Hsieh, J.Y., et al. (2020). YAP-mediated mechanotransduction tunes the macrophage inflammatory response. Sci Adv 6. 10.1126/sciadv.abb8471.

33. Scott, K.E., Fraley, S.I., and Rangamani, P. (2021). A spatial model of YAP/TAZ signaling reveals how stiffness, dimensionality, and shape contribute to emergent outcomes. Proc Natl Acad Sci U S A 118. 10.1073/pnas.2021571118.

34. Takahashi, S., Leiss, M., Moser, M., Ohashi, T., Kitao, T., Heckmann, D., Pfeifer, A., Kessler, H., Takagi, J., Erickson, H.P., et al. (2007). The RGD motif in fibronectin is essential for development but dispensable for fibril assembly. Journal of Cell Biology 178, 167–178. 10.1083/JCB.200703021.

35. Perugorria, M.J., Olaizola, P., Labiano, I., Esparza-Baquer, A., Marzioni, M., Marin, J.J.G., Bujanda, L., and Banales, J.M. (2018). Wnt–β-catenin signalling in liver development, health and disease. Nature Reviews Gastroenterology & Hepatology 2018 16:2 16, 121–136. 10.1038/s41575-018-0075-9.

36. Rausch, V., and Hansen, C.G. (2020). The Hippo Pathway, YAP/TAZ, and the Plasma Membrane. Trends Cell Biol 30, 32–48. 10.1016/J.TCB.2019.10.005.

37. Gu, B., Bradshaw, B., Zhu, M., Sun, Y., Hopyan, S., and Rossant, J. (2022). Live imaging YAP signalling in mouse embryo development. Open Biol 12. 10.1098/rsob.210335.

38. Belteki, G., Haigh, J., Kabacs, N., Haigh, K., Sison, K., Costantini, F., Whitsett, J., Quaggin, S.E., and Nagy, A. (2005). Conditional and inducible transgene expression in mice through the combinatorial use of Cre-mediated recombination and tetracycline induction. Nucleic Acids Res 33, 1–10. 10.1093/nar/gni051.

39. Muzumdar, M.D., Tasic, B., Miyamichi, K., Li, N., and Luo, L. (2007). A global double-fluorescent cre reporter mouse. Genesis 45. 10.1002/dvg.20335.

40. Perantoni, A.O., Timofeeva, O., Naillat, F., Richman, C., Pajni-Underwood, S., Wilson, C., Vainio, S., Dove, L.F., and Lewandoski, M. (2005). Inactivation of FGF8 in early mesoderm reveals an essential role in kidney development. Development. 10.1242/dev.01945.

41. Reginensi, A., Scott, R.P., Gregorieff, A., Bagherie-Lachidan, M., Chung, C., Lim, D.S., Pawson, T., Wrana, J., and McNeill, H. (2013). Yap- and Cdc42-Dependent Nephrogenesis and Morphogenesis during Mouse Kidney Development. PLoS Genet 9. 10.1371/journal.pgen.1003380.

42. Yamaguchi, T.P., Bradley, A., McMahon, A.P., and Jones, S. (1999). A Wnt5a pathway underlies outgrowth of multiple structures in the vertebrate embryo. Development.

43. Ferrer-Vaquer, A., Piliszek, A., Tian, G., Aho, R.J., Dufort, D., and Hadjantonakis, A.K. (2010). A sensitive and bright single-cell resolution live imaging reporter of Wnt/-catenin signaling in the mouse. BMC Dev Biol 10. 10.1186/1471-213X-10-121.

44. Ohashi, T., Kiehart, D.P., and Erickson, H.P. (1999). Dynamics and elasticity of the fibronectin matrix in living cell culture visualized by fibronectin-green fluorescent protein. Proc Natl Acad Sci U S A 96. 10.1073/pnas.96.5.2153.

45. Chu, V.T., Weber, T., Graf, R., Sommermann, T., Petsch, K., Sack, U., Volchkov, P., Rajewsky, K., and Kühn, R. (2016). Efficient generation of Rosa26 knock-in mice using CRISPR/Cas9 in C57BL/6 zygotes. BMC Biotechnol 16. 10.1186/s12896-016-0234-4.

46. Gu, B., Posfai, E., Gertsenstein, M., and Rossant, J. (2020). Efficient Generation of Large-Fragment Knock-In Mouse Models Using 2-Cell (2C)-Homologous Recombination (HR)-CRISPR. Curr Protoc Mouse Biol 10. 10.1002/cpmo.67.

47. Gu, B., Gertsenstein, M., and Posfai, E. (2020). Generation of Large Fragment Knock-In Mouse Models by Microinjecting into 2-Cell Stage Embryos. In Methods in Molecular Biology 10.1007/978-1-4939-9837-1_7.

48. Fenelon, K.D., Thomas, E., Samani, M., Zhu, M., Tao, H., Sun, Y., McNeill, H., and Hopyan, S. (2022). Transgenic force sensors and software to measure force transmission across the mammalian nuclear envelope in vivo. Biol Open 11. 10.1242/BIO.059656/281166.

49. Wong, M.D., Dazai, J., Walls, J.R., Gale, N.W., and Henkelman, R.M. (2013). Design and Implementation of a Custom Built Optical Projection Tomography System. PLoS One 8. 10.1371/journal.pone.0073491.

