## Supplemental Figures for "Durotaxis bridges phase transition as a function of tissue stiffness *in vivo*"

Figure S1

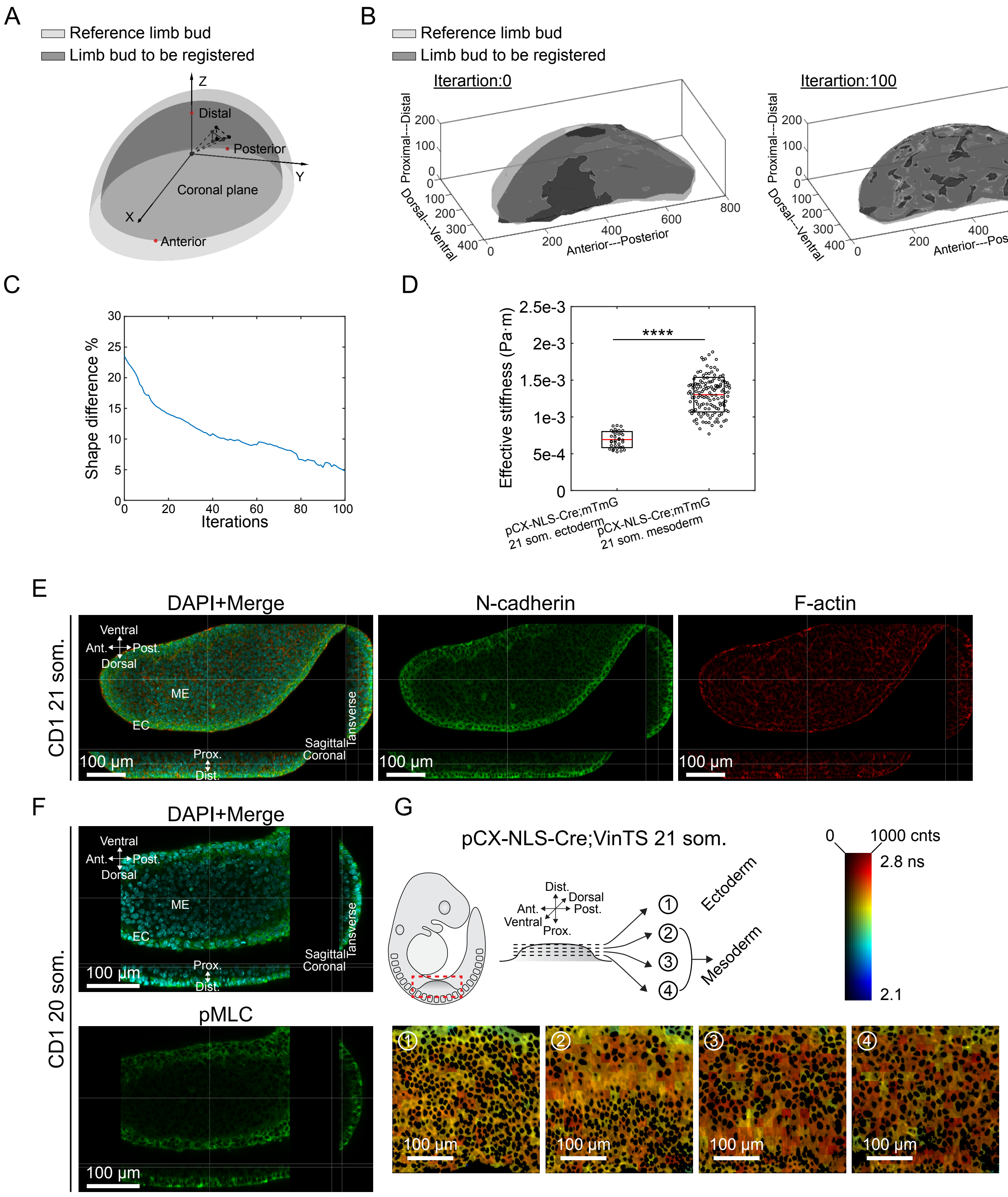


**Three-dimensional registration of limb bud shapes.** (A) Schematic illustrating the 3D limb bud shape registration. (B) Representative 3D limb bud shape registration after 100 iterations. (C) Representative limb bud shape difference percentage at different iterations of registration. (D) Effective stiffness values of data shown in Figure 1B (two-tailed unpaired Student's t-test, ****P < 0.0001). **Cell migration is unlikely to be driven by cell sorting or biased force generation.** (E) Confocal sections of 21 som. CD1 embryos at forelimb region visualising DAPI (cyan), anti-N-cad-herin antibody (green) and F-actin (red). (F) Confocal sections of 20 som. CD1 embryos at forelimb region visualising DAPI (cyan) and pMLC (green). (G) Three-dimensional rendering of the 21 som. stage pCX-NLS-Cre;VinTS limb buds vinculin tension sensor lifetime map.

Figure S2

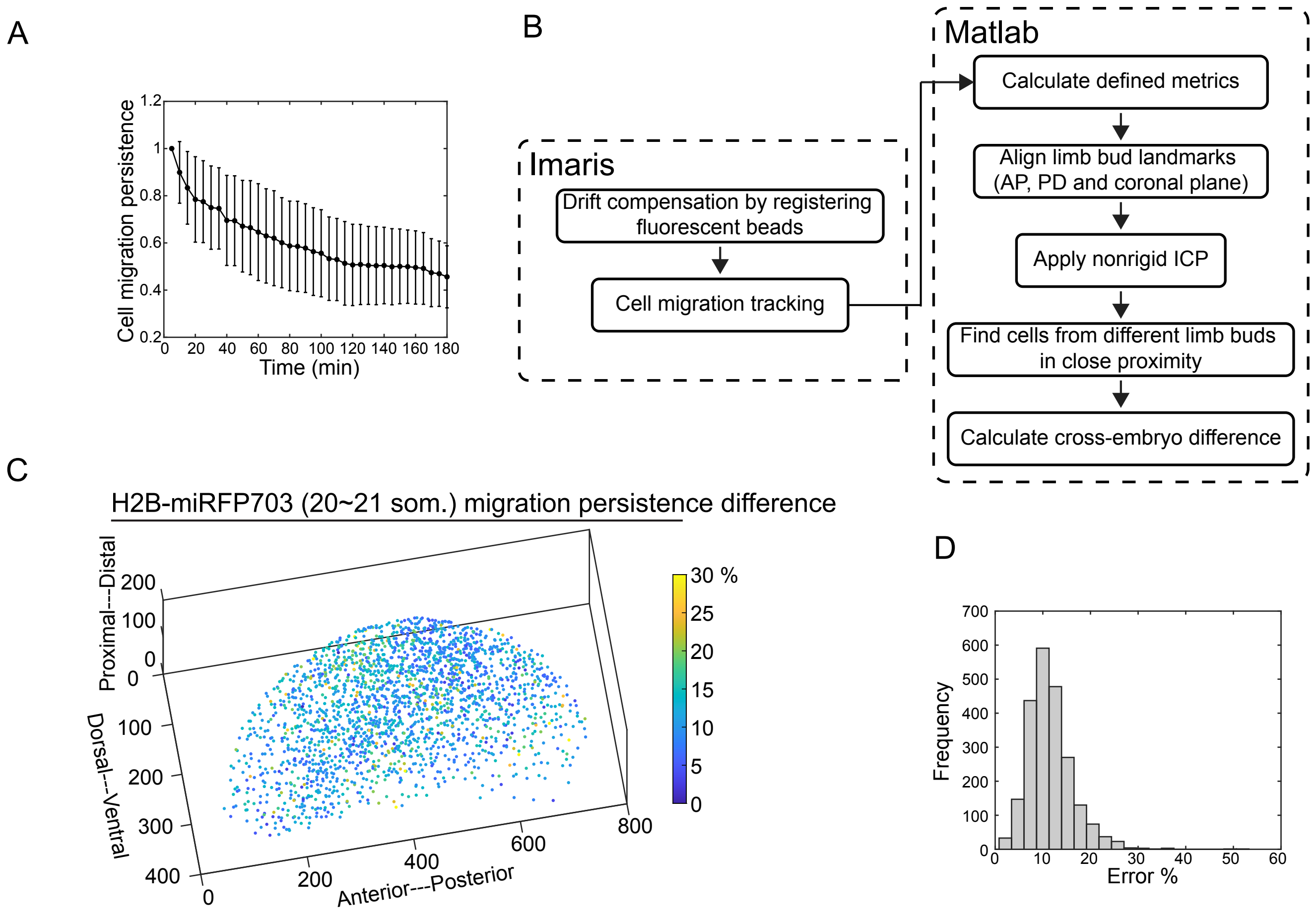


**Cell migration persistence analysis.** (A) H2B-miRFP703 21 som. limb bud average mesodermal cell migration persistence at different timed points. (B) Flow chart of calculating cross-embryo difference in cell migration persistence. (C) H2B-miRFP703 20~21 som. limb buds cell migration persistence difference. (D) Histogram representing the error percentage of cross-embryo difference in cell migration persistence shown in (C). Error bars indicate s.d..

Figure S3

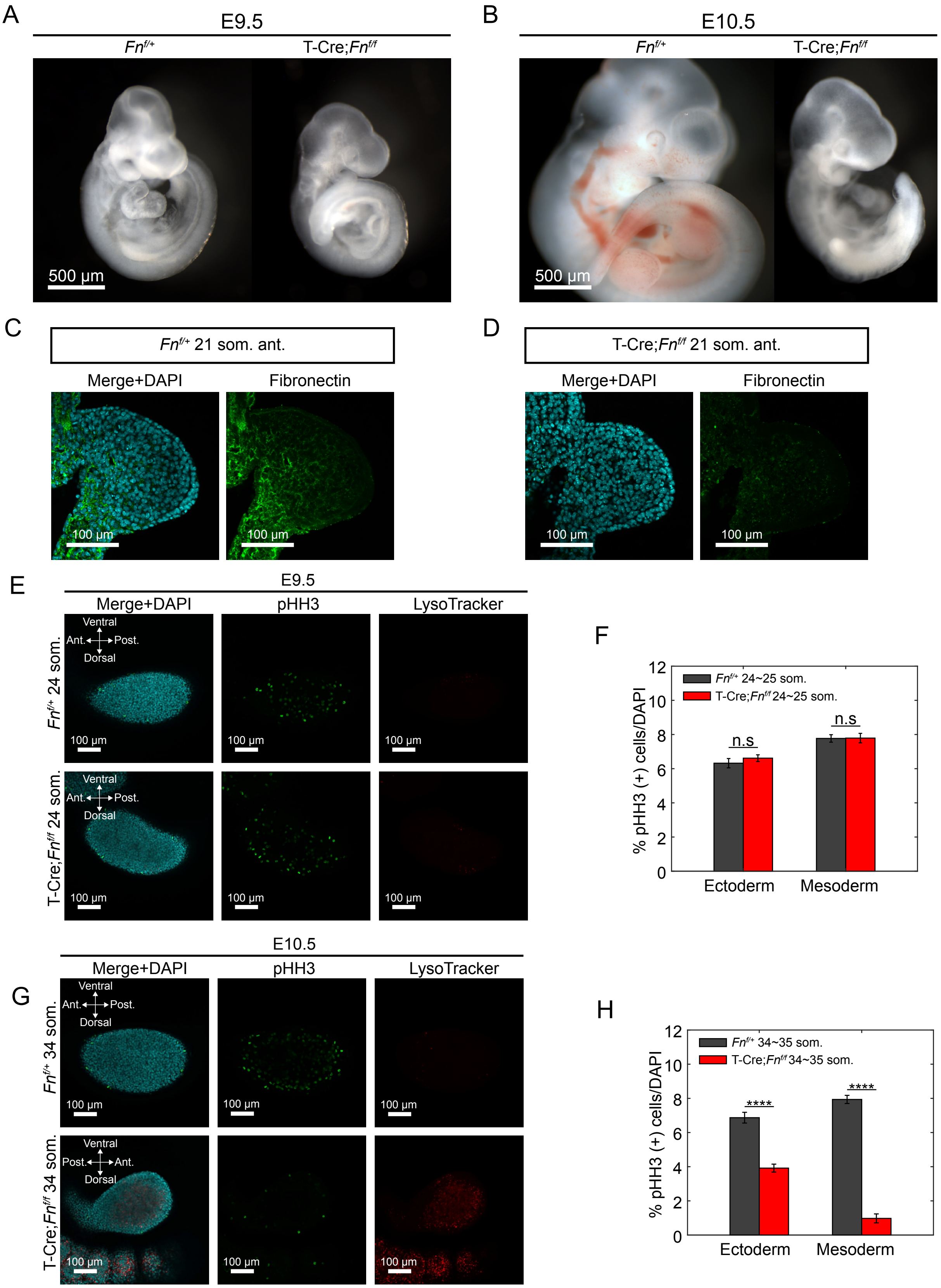


**Conditional knock-out of fibronectin.** (A and B) Morphology of *Fn*$^{f/+}$ and T-Cre;*Fn*$^{f/f}$ embryos at E9.5 (A) and E10.5 (B). (C and D) Transverse sections of 21 som. *Fn*$^{f/+}$ (C) and T-Cre;*Fn*$^{f/f}$ (D) forelimb at anterior region. Sections were stained with DAPI (cyan) and fibronectin antibody (green). (E) Confocal sections of 24 som. *Fn*$^{f/+}$ and T-Cre;*Fn*$^{f/f}$ limb buds visualising DAPI (cyan), anti-pHH3 antibody (green) and LysoTracker (red). (F) Histogram representing percentage of pHH3-positive cells in the 24~25 som. T-Cre;*Fn*$^{f/f}$ limb bud (n=3) in both ectoderm and mesoderm (two-tailed unpaired Student's t-test, ****P<0.0001). (E) Confocal sections of 34 som. *Fn*$^{f/+}$ and T-Cre;*Fn*$^{f/f}$ limb buds visualising DAPI (cyan), anti-pHH3 antibody (green) and LysoTracker (red). (G) Histogram representing percentage of pHH3-positive cells in the 34~35 som. T-Cre;*Fn*$^{f/f}$ limb bud (n=3) in both ectoderm and mesoderm (two-tailed unpaired Student's t-test). n.s, not significant.

Figure S4

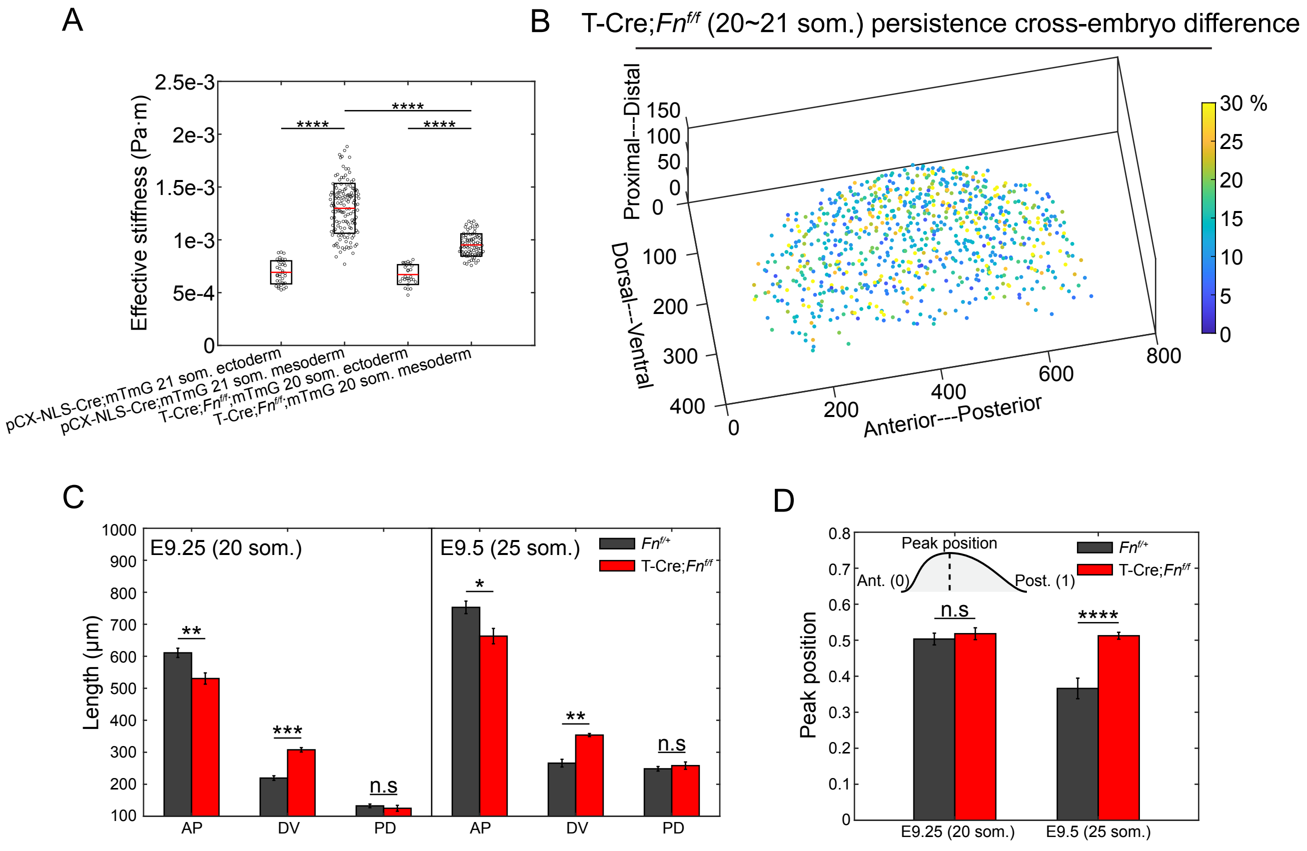


**Loss of fibronectin downregulates tissue stiffness and leads to a broadly rearranging state.** (A) Effective stiffness values of data shown in in Figure 1B and Figure 2A (two-tailed unpaired Student's t-test, ****P < 0.0001). (B) T-Cre;*Fn*$^{f/f}$ 20~21 som. limb buds cell migration persistence difference.(C) Histogram representing the AP, DV and PD axes length of 20 and 25 som. *Fn*$^{f/+}$ and T-Cre;*Fn*$^{f/f}$ limb buds (two-tailed unpaired Student's t-test, *P<0.05, **P<0.01, ***P<0.001, n = 3 embryos for each condition). (D) Histogram representing the distally biased peak location in the AP axis of 20 and 25 som. *Fn*$^{f/+}$ and T-Cre;*Fn*$^{f/f}$ limb buds (two-tailed unpaired Student's t-test, ****P<0.0001). n.s, not significant. Error bars indicate s.d..

Figure S5

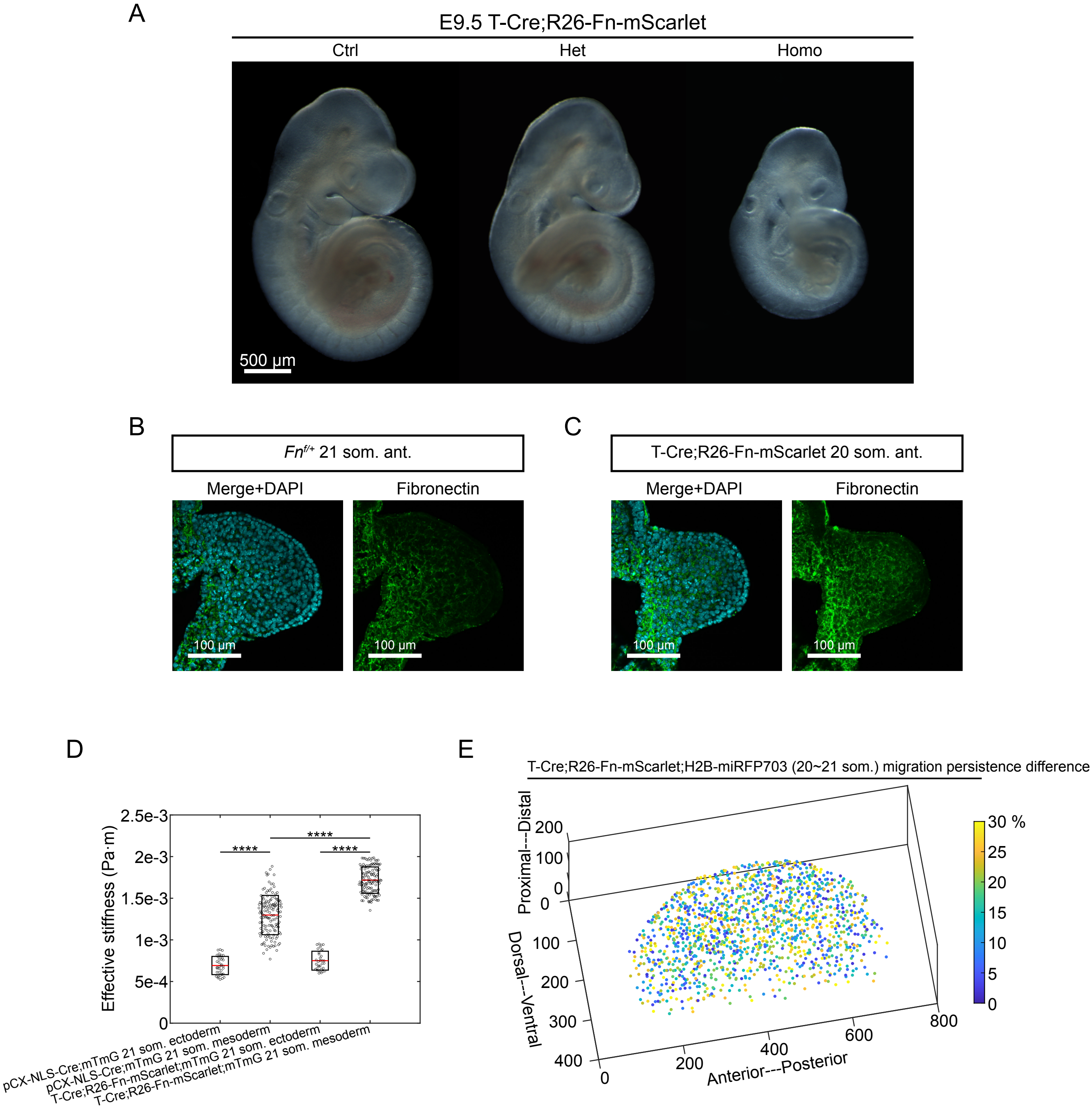


**Conditional overexpression of fibronectin upregulates tissue stiffness and leads to a broadly caged state.** (A) Morphology of T-Cre;R26-Fn-mScarlet embryos at E9.5. (B and C) Transverse sections of 21 som. *Fn*$^{f/+}$ (B) and 20 som. T-Cre;R26-Fn-mScarlet (C) forelimb at anterior region. Sections were stained with DAPI (cyan) and fibronectin antibody (green). (D) Effective stiffness values of data shown in Figure 1B and Figure 3A (two-tailed unpaired Student's t-test, ****$P < 0.0001$). (E) T-Cre;R26-Fn-mScarlet;H2B-miRFP703 20~21 som. limb buds cell migration persistence difference.

### Figure S6

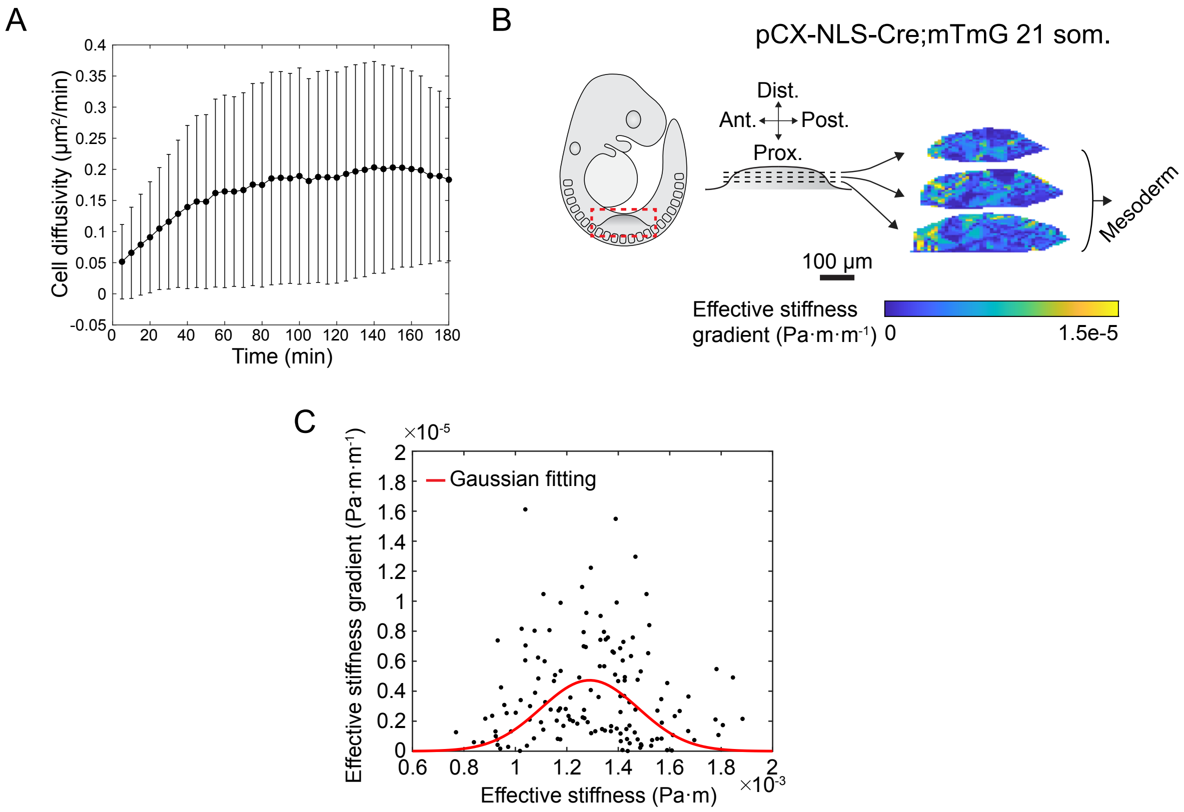


**Modified Landau phase diagram.** (A) Average cell diffusivity of H2B-miRFP703 20 som. limb bud over 3 h imaging period. (B) Three-dimensional rendering of the 21 som. stage pCX-NLS-Cre;mT-mG limb buds mesodermal tissue stiffness gradient map (n=5). (C) Gaussian fitting of stiffness gradient vs. stiffness.

Figure S7

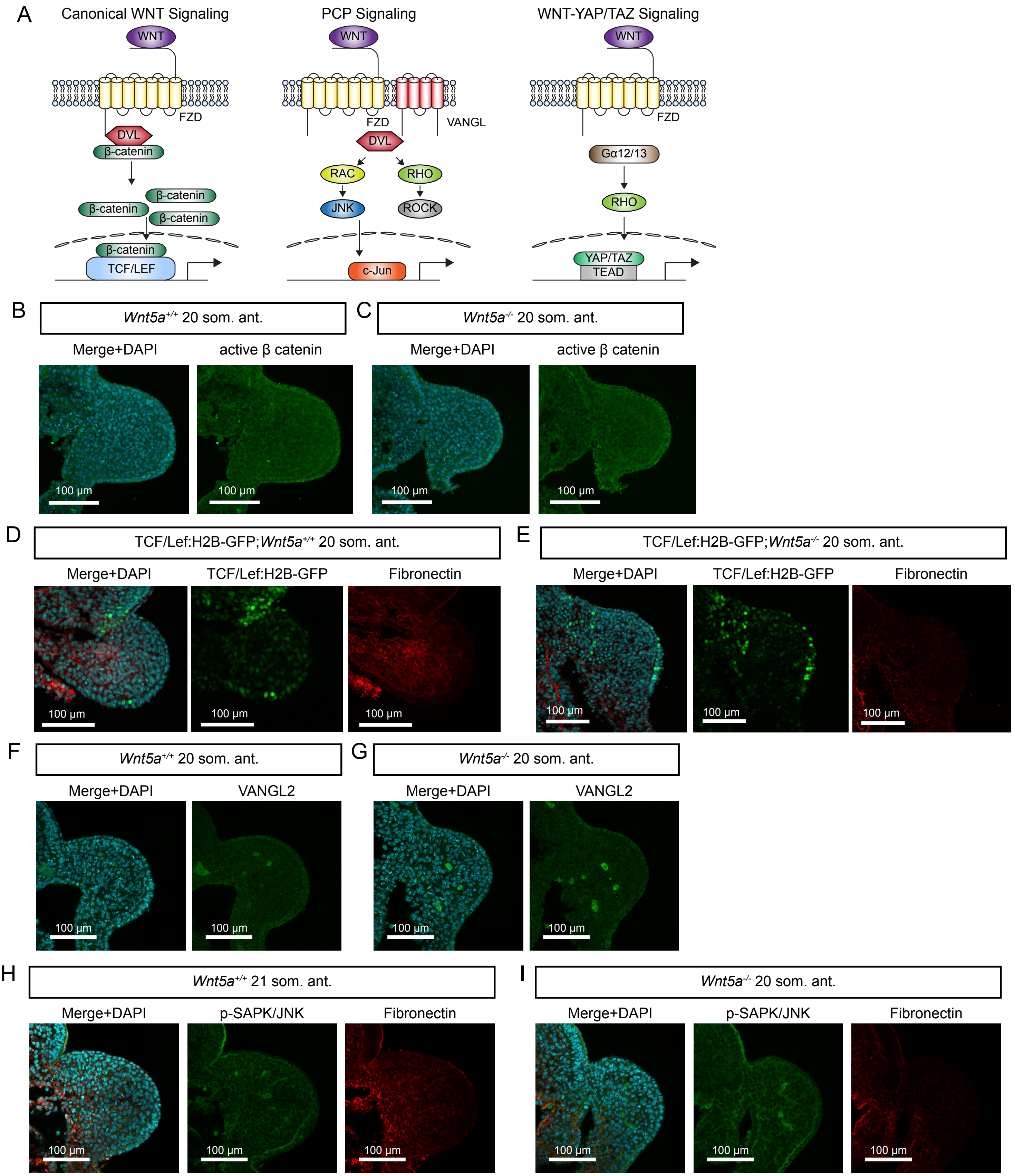


**Immunostaining against WNT pathway components.** (A) Schematic representing WNT signalling pathways including canonical, PCP and WNT-YAP/TAZ. (B and C) Transverse sections of 20 som. *Wnt5a*$^{+/-}$ (B) and *Wnt5a*$^{-/-}$ (C) forelimb at anterior region. Sections were stained with DAPI (cyan) and anti-beta catenin antibody (green). (D and E) Transverse sections of 20 som. TCF/Lef:H2B-GFP;*Wnt5a*$^{+/-}$ (D) and TCF/Lef:H2B-GFP;*Wnt5a*$^{-/-}$ (E) forelimb at anterior region. Sections were stained with DAPI (cyan) and anti-fibronectin antibody (red). (F and G) Transverse sections of 20 som. *Wnt5a*$^{+/-}$ (F) and *Wnt5a*$^{-/-}$ (G) forelimb at anterior region. Sections were stained with DAPI (cyan) anti-VANGL2 antibody (green). (H and I) Transverse sections of 20 som. *Wnt5a*$^{+/-}$ (H) and *Wnt5a*$^{-/-}$ (I) forelimb at anterior region. Sections were stained with DAPI (cyan), anti-p-SAPK antibody (green) and anti-fibronectin antibody (red).

Figure S8

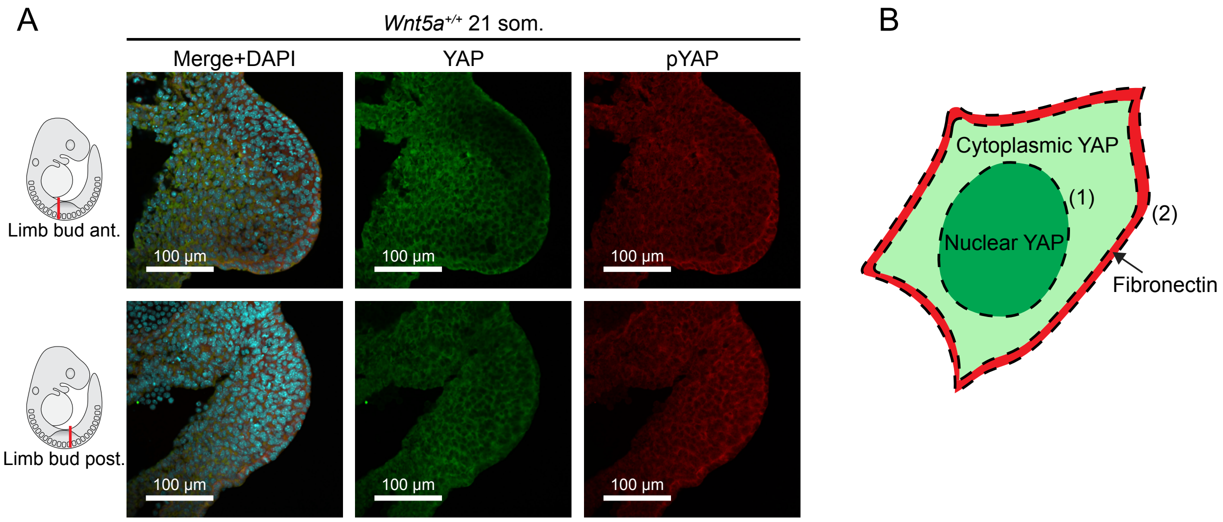


**Immunostaining against YAP.** (A) Transverse sections of 21 som. *Wnt5a*$^{+/+}$ embryos at anterior and posterior forelimb regions. Sections were stained with DAPI (cyan) anti-YAP (green) and anti-phospho-YAP antibody (red). (B) Schematic illustrating analysis method of FN and YAP nuclear/cytoplasmic ratio colocalisation.
